# Single-cell landscapes of primary glioblastomas and matched organoids and cell lines reveal variable retention of inter- and intra-tumor heterogeneity

**DOI:** 10.1101/2021.04.24.441206

**Authors:** VG LeBlanc, DL Trinh, S Aslanpour, M Hughes, D Livingstone, MD Blough, JG Cairncross, JA Chan, JJ Kelly, MA Marra

## Abstract

Glioblastomas (GBMs) are aggressive primary malignant brain tumors characterized by extensive levels of inter- and intra-tumor genetic and phenotypic heterogeneity. Patient-derived organoids (PDOs) have recently emerged as useful models to study such heterogeneity. Here, we present bulk exome as well as single-cell genome and transcriptome profiles of primary *IDH* wild type GBMs from ten patients, including two recurrent tumors, as well as PDOs and brain tumor-initiating cell (BTIC) lines derived from these patients. We find that PDOs are genetically similar to and variably retain gene expression characteristics of their parent tumors. At the phenotypic level, PDOs appear to exhibit similar levels of transcriptional heterogeneity as their parent tumors, whereas BTIC lines tend to be enriched for cells in a more uniform transcriptional state. The datasets introduced here will provide a valuable resource to help guide experiments using GBM-derived organoids, especially in the context of studying cellular heterogeneity.

## Introduction

Malignant gliomas account for approximately 80% of primary malignant brain tumors. Glioblastoma (GBM) is the most common and aggressive subtype of glioma, with a five-year survival rate in adults of only 4.3% (Ostrom et al., 2018). Although histopathologically defined as grade IV astrocytomas, GBMs display extensive genomic, epigenomic, metabolic, and microenvironmental heterogeneity both between patients (inter-tumor) and within individual tumors (intra-tumor) (Vartanian et al., 2014). This heterogeneity has confounded our understanding of the cellular and molecular underpinnings of GBM and has complicated the development of desperately needed therapies. To address the lack of detailed data describing GBM molecular heterogeneity, single-cell sequencing technologies have been deployed to measure genomic (Francis et al., 2014; Guilhamon et al., 2021; Wang et al., 2019) and transcriptomic (Bhaduri et al., 2020; Castellan et al., 2021; Couturier et al., 2020; Darmanis et al., 2017; Garofano et al., 2021; Lee et al., 2017; Muller et al., 2017; Neftel et al., 2019; Patel et al., 2014; Richards et al., 2021; Venteicher et al., 2017; Wang et al., 2019) heterogeneity. Such analyses have facilitated deeper explorations of inter- and intra-tumor heterogeneity than previously possible, and are poised to inform on the contributions made by molecular and cellular heterogeneity to key disease features such as drug resistance and tumor recurrence (Dagogo-Jack and Shaw, 2017). Recently, Neftel *et al*. (2019) showed that transcriptional signatures of intra-tumor heterogeneity in GBM converged on cell cycle state or one of four cell states reminiscent of developmental stages observed in the brain (neural progenitor-, oligodendrocyte progenitor-, astrocytic-, and mesenchymal-like). Despite associations between specific mutational drivers and an enrichment of these cell states, the translational potential of such observations will rely, at least in part, on the use of experimental models that can reproducibly and tractably recapitulate these observations.

Organoids have emerged as useful models capable of recapitulating biological features of both normal and disease tissues, in part because they appear to allow for the propagation of the heterogeneity of cell types encountered in tissues (Lou and Leung, 2017; Weeber et al., 2017). For example, patient-derived GBM organoids (PDOs) were shown to retain cellular heterogeneity reminiscent of that observed in primary tumors, such as hypoxic cells near the tumor core and cycling stem-like cells near the invasive rim (Hubert et al., 2016; Jacob et al., 2020). Interestingly, Hubert *et al*. (2016) also showed that xenografts derived from organoids and those derived from neurospheres differed in their latency and histology, with the former being more representative of the original tumor and displaying behavior typical of GBM such as infiltration into the surrounding brain. Similarly, others have implanted neurospheres into cerebral organoids, creating GBM models that allow for host-tumor cell interactions that closely resemble those observed *in vivo* (cerebral organoid gliomas, GLICOs) (Linkous et al., 2019; Ogawa et al., 2018). Early studies thus indicated that PDOs and GLICOs may present an improved *in vitro* model to study GBM, especially in the context of inter- and intra-tumor heterogeneity and the effects of these on disease progression and treatment response. The extent to which organoid models recapitulate the genetic and molecular heterogeneity seen in primary GBM needs to be carefully examined at the single-cell level to provide a thorough understanding of their cellular makeup and their potential to address outstanding questions in the field.

Here, we report the generation and analysis of bulk whole-exome, >8,000 single-cell genome, and >75,000 single-cell transcriptome profiles derived from primary GBMs, matched PDOs, and brain tumor-initiating cell (BTIC) lines. We present data from a total of 10 tumors, with each tumor represented by samples from two non-adjacent tumor locations selected from the primary GBM resection. PDOs were derived from each of these non-adjacent regions, and replicate PDOs (up to three per region) were profiled when they were available. To compare transcriptome-level single-cell heterogeneity between organoids and BTIC lines derived from the same tumor sample, we also derived and profiled BTIC lines from one or both tumor regions of five GBMs.

## Results

### A novel approach for developing GBM-derived organoids

To explore the extent to which novel and established model systems recapitulate the inter- and intra-tumor heterogeneity previously reported in GBM and described above, we first obtained primary tissue samples from two spatially distinct regions of primary IDH wild-type GBMs from 10 patients and then created PDOs from these samples and BTIC lines from a subset of them (Supplementary Table 1). For two of the patients (JK136 and JK142), we also obtained samples from a single region of tumors that had recurred in these patients and created PDOs from these samples. To obtain PDOs, we developed a protocol to create organoids from intact (*i.e.* non-dissociated) tumor sections embedded in Matrigel (Figure 1A; Methods). Briefly, freshly resected tumor tissue from the enhancing margin of each tumor was obtained and cut into small sections (∼1 mm^3^), then rinsed and placed into prepared Matrigel forms. After being allowed to solidify at room temperature for 10 mins, additional Matrigel was added to seal the tumor section within the resulting Matrigel sphere. Each organoid was then cultured in 24-well plates containing neural stem cell media supplemented with EGF, FGF-2, and heparin sulfate. Organoid growth was typically observed after 24-48 hrs, and the organoids were maintained with media changes as necessary until they grew out into the surrounding Matrigel (range 17-55 days), at which point they were either cryopreserved or sectioned. In some cases, we observed cells escaping the Matrigel and creating neurospheres within the well; when this occurred, these cells were further propagated as BTIC lines (Methods).

**Figure 1:**
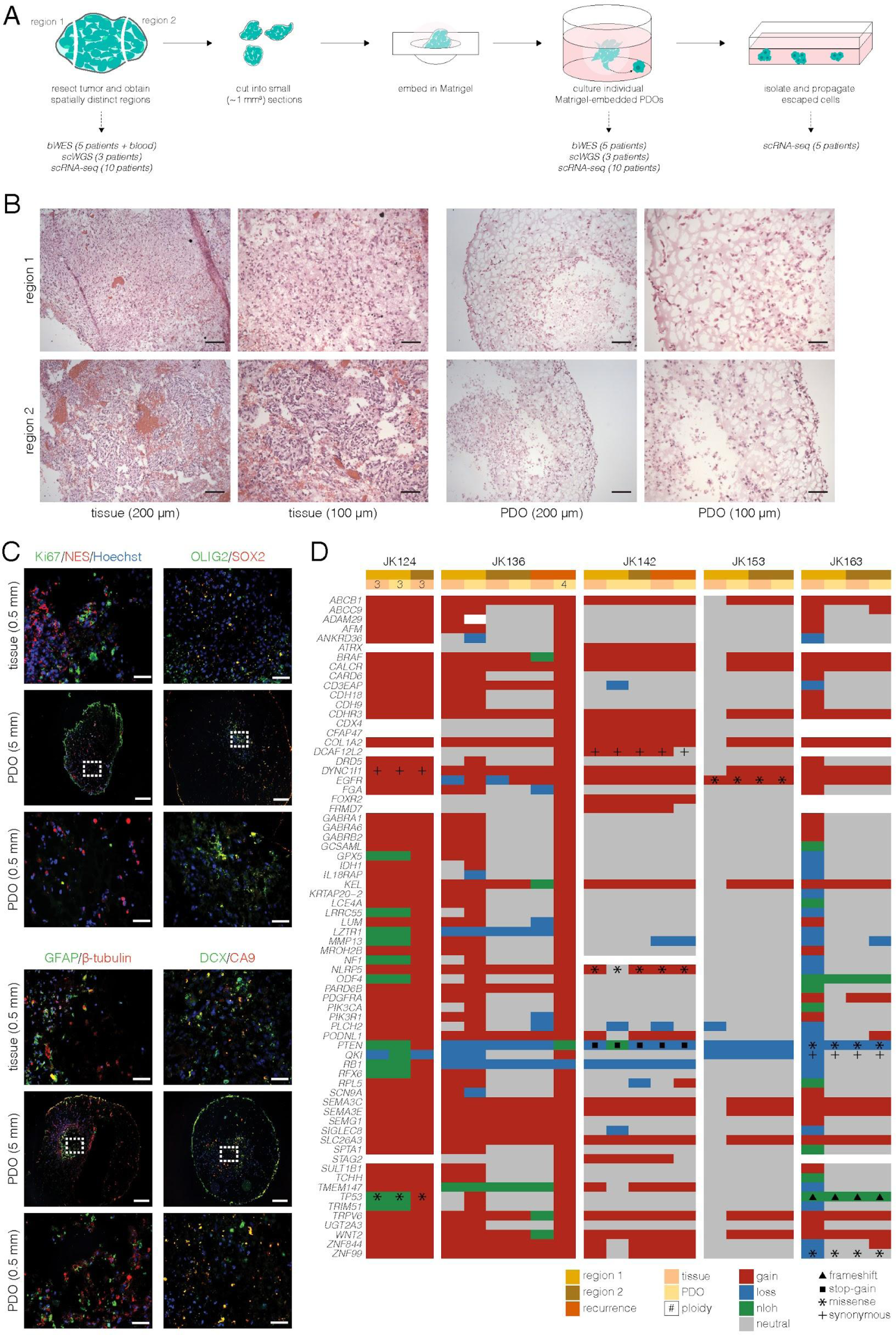
Generation and characterization of patient-derived organoids. **A** Schematic of the process used to generate PDOs and BTIC lines (see Methods for more details). Two spatially distinct regions were obtained from opposite ends of each resected tumor — the naming (region 1 vs region 2) does not indicate any consistent feature and is thus not comparable between tumors/patients. Small intact sections were obtained from each region and embedded in Matrigel spheres, which were then cultured (one sphere/well). In some cases, cells escaped the Matrigel sphere and seeded new cultures in the well; these were isolated and propagated as BTIC lines. Original tissue sections, PDOs, and BTIC lines were all profiled for this study, with details indicated below the schematic. **B** Representative image showing H&E staining of matched tissue/PDO pairs (JK125). Scale bar lengths are indicated below the images. **C** Representative immunofluorescence images showing similar expression patterns of select markers in matched tissue/PDO pairs. White boxes on PDOs indicate the area shown in the zoomed-in images below. Hoescht staining is shown in blue in all images. Ki67/NES: JK156 reg1; OLIG2/SOX2: JK153 reg2; GFAP/β-tubulin: JK152 reg1; DCX/CA9: JK125 reg2. Scale bar lengths are indicated on the images’ left. **D** Copy number alterations and SNVs/indels identified in GBM genes across samples. Ploidy is indicated for polyploid samples (others are diploid). Full variant information can be found in Supplementary Table 2. nloh: copy number neutral loss of heterozygosity.

To determine whether the PDOs resembled their parental tumors at the histological level, we performed hematoxylin and eosin (H&E) staining on matched tissues and PDOs (example shown in Figure 1B). A neuropathologist confirmed that the PDOs retain features of their parental tumors that are typical to GBM, such as nuclear atypia, mitotic figures, and variable cellular morphology. The PDOs also all show some degree of invasion from the cellular mass into the adjacent matrix. Additionally, immunofluorescence analyses showed that PDOs had typical GBM features, such as cycling cells (Ki67+), cells expressing stem- or progenitor-like markers (NES, SOX2, OLIG2, DCX) and glial markers (GFAP), as well as hypoxic cells (CA9+; Figure 1C). Together, these results indicate that PDOs retain key features of their parent tumors.

### PDOs retain genomic characteristics of their parent tumor sample

To analyze the extent to which PDOs could preserve the mutational heterogeneity present in parent tumors, we generated bulk whole-exome sequencing (bWES) libraries from 29 samples from five patients (Supplementary Table 1) at a median coverage depth of ∼260X. We called somatic variants, both copy number and single nucleotide variants (SNVs) and small insertions/deletions (indels), using matched blood as normal comparators (Methods). Two samples, namely the JK124 reg2 PDO and the JK142 reg2 tissue sets, had low inferred tumor content (discussed in more detail below); however, all tumors with sufficiently high tumor content to call somatic alterations had multiple alterations in genes that have been implicated in GBM (Brennan et al., 2013), and in most cases variants were commonly found in different regions of the same tumor and in matched tissue/PDO pairs (Figure 1D).

To quantify similarities between tissues and PDOs, we first used PyClone (Roth et al., 2014) to cluster variants from all samples belonging to each patient. Comparing PyClone cluster prevalences across different sample pairs, we found similar or better concordance between tissue/PDO pairs than between tissue samples taken from opposite regions of the same tumor (Supplementary Figure 1). This suggests that, even in cases where a tumor has high regional variant heterogeneity, PDOs largely retain variants at comparable frequencies found in the tissue from which they were derived. We then used CITUP (Malikic et al., 2015) to determine the phylogenetic relationships between these variant clusters and to identify the cellular prevalence of each clone across samples (*i.e.* the proportion of malignant cells belonging to each clone). Overall, these proportions were again comparable between PDOs and matched tissue samples, indicating that the PDOs largely retain the hierarchical structures observed in their parental tissues (Figure 2). Even when proportions of individual clones differed, matched tissues and PDOs tended to be composed of clones from the same branch(es) (*e.g.* clones 9 and 10 in JK142 reg1, Figure 2F), indicating the possibility of some evolutionary growth in the PDOs or possibly sampling differences. In both patients with recurrent samples, clones that were unique to the recurrent samples (clone 4 in JK136, Figure 2C; clones 4 and 5 in JK142, Figure 2F) were notably present in both the tissue and PDO samples. Together, these results indicated that the PDOs largely retained the clonal structures observed in their parent tissues.

**Figure 2:**
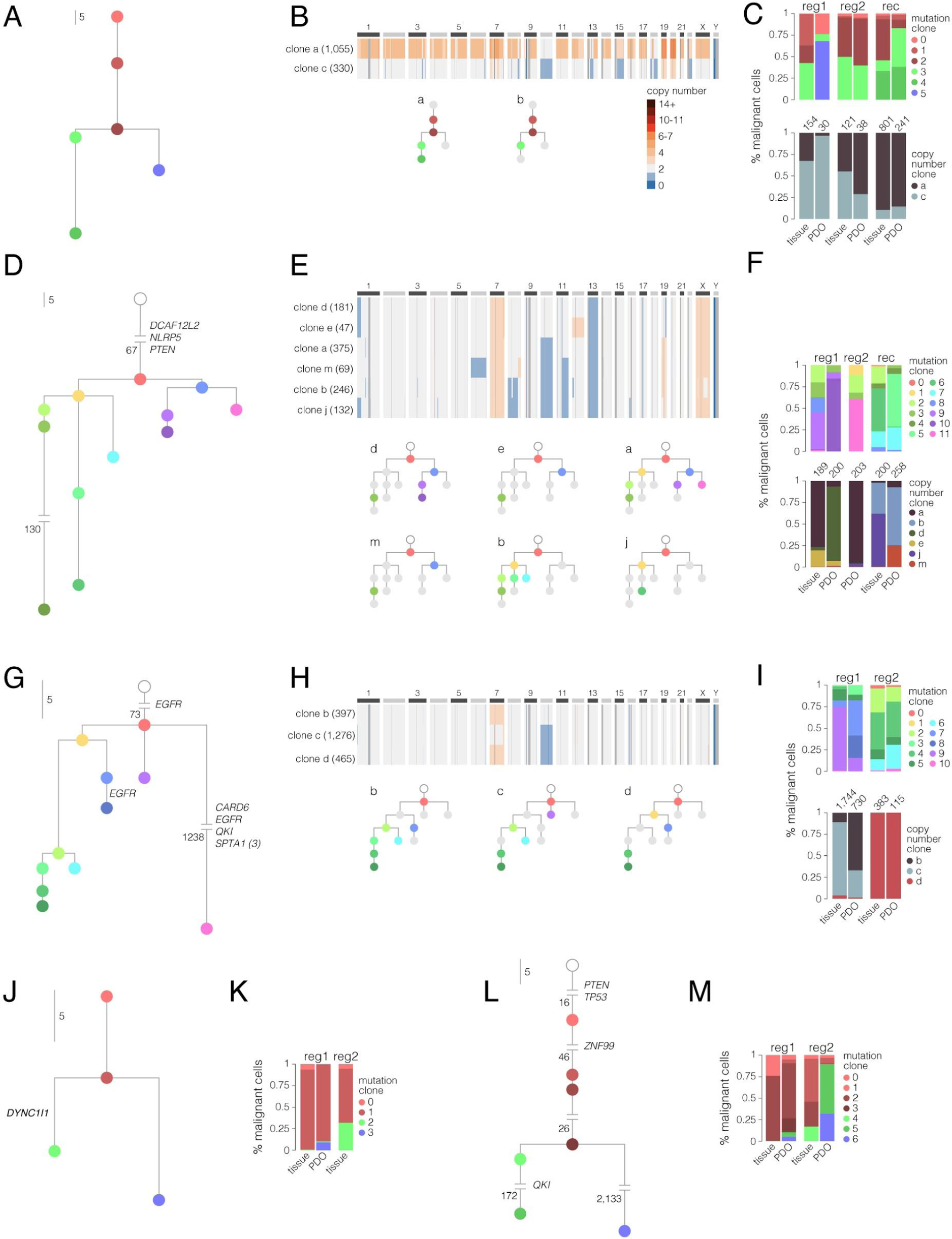
PDOs retain genetic characteristics of their parent tumors. Clonal hierarchies identified in samples from JK136 (**A-C**), JK142 (**D-F**), JK153 (**G-I**), JK124 (**J-K**), and JK163 (**L-M**). **A**,**D**,**G**,**J**,**L** Trees were derived using CITUP with mutation clusters obtained using PyClone (bWES data). Edge lengths are proportional to the number of mutations gained in each clone (node), except where edges are broken for display purposes; in such cases, the number of mutations is indicated for broken edges. GBM-associated genes (Brennan et al., 2013) carrying mutations are indicated when present. **B**,**E**,**H** Median copy number profiles of malignant copy number clones identified using scWGS data. The number of cells in each clone is indicated in brackets, and the copy number color legend is shown in panel B. Mutation clones on the accompanying tree schematics are colored if the copy number clone carries evidence of >10% variants from the mutation clone (Methods). **C**,**F**,**I**,**K**,**M** Proportion of malignant cells assigned to each mutation clone (all tumors) and each copy number clone (JK136, JK142, and JK153; bottom, C,F,I). The number of malignant cells identified by scWGS is shown for each sample.

While tissues and matched PDOs tended to carry a comparable number of variants (Supplementary Figure 2), JK163 appeared to present a unique case in which both PDOs displayed much higher numbers of distinct variants than their matched tissues: 1,538 variants were identified in the reg1 PDO (14-fold higher than the matched tissue) and 22,248 variants were identified in the reg2 PDO (227-fold higher than the matched tissue). Both JK163 PDOs thus appeared to have gained a hypermutation phenotype, with mutation burdens of 38 and 550 mutations per megabase, respectively. To further investigate the possible cause of these events, we calculated exposure to single-base substitution signatures recently defined by the Pan-Cancer Analysis of Whole Genomes (PCAWG) consortium (Alexandrov et al., 2020). Both tissue samples mostly showed exposure to signature SBS1, which is related to age; conversely, both PDO samples showed highest exposure to signature SBS39, whose aetiology is unknown (Supplementary Figure 2). Although mutation signature analysis was less reliable in other samples with fewer mutations, none of the other PDO or tissue samples showed an enrichment of SBS39 (Supplementary Table 3), indicating that this observation is unlikely to be driven exclusively by the PDO culture conditions. Mutations in mismatch repair (MMR) genes are relatively common in GBM and often arise as a resistance mechanism to temozolomide treatment (Touat et al., 2020). Although a stop-gain variant was identified in *MSH6* in this patient, its allelic fraction was highest in the tissue sample from reg1 (2.2%) and was in fact lower in PDOs (0.9% in both regions), which is inconsistent with the notion that an MMR-deficient clone evolved to dominance in the PDOs leading to high mutational burdens. No variants were found in other MMR genes. The foundational clone identified in this patient’s samples did however carry deleterious mutations in both *TP53* and *PTEN*, and was notably the only such case in our dataset. The PDOs were also the second most rapidly growing of all the samples analyzed here (harvested at day 19, range 17-55 days), which could imply a relationship between rapid growth and high mutation burden in some contexts.

### Copy number-based clonal populations in parent tissues are retained in PDOs

To directly observe clonal properties rather than infer them from bulk sequencing data, we further identified and compared subclonal populations in tissues and PDOs from three patients using single-cell whole-genome sequencing (scWGS) to identify copy number alterations (Supplementary Table 1). We sequenced 115-1,779 cells per sample (median 384), reaching a median of 665,324 reads per cell across samples. For each patient, we identified clonal populations defined by clusters of cells with similar copy number profiles (Figure 2B/E/H; Methods). Notably, both the bWES and scWGS datasets indicated relatively rare malignant cell populations in the JK142 region 2 tissue sample (tumor content identified as 0.28% for the bWES data, and 8/485 cells [1.6%] identified as malignant in the scWGS data). In the JK153 samples, the malignant copy number clones were consistent with a polyclonal origin, as clone b contained a chromosome 7 amplification, clone c contained a chromosome 10 deletion, and clone d contained both events (Figure 2H). However, allelic fractions of variants identified as heterozygous in the germline sample (from the bWES data) indicated that there was loss of heterozygosity (LOH) affecting chromosome 10 in all three clones (Supplementary Figure 2). These data may thus indicate that clone b was derived from a clone d cell that experienced a duplication of its remaining copy of chromosome 10. Across patients and as observed for the mutation clones, comparable proportions of malignant cells were assigned to distinct copy number clones in matched tissues and PDOs, indicating that the PDOs also retained copy number-based clonal characteristics observed in their parent tumors.

We then sought to integrate the single-cell copy number data with our bWES data. To do so, we pooled the sequencing reads from all the cells assigned to each copy number clone, and then screened these for the existence of mutations we had called in the bWES data (Methods). Most copy number clones contained mutations from more than one mutation clone, consistent with the notion that copy number events occur less frequently than SNVs and INDELs (Figures 2B/E/H). Additionally, recurrent samples (both tissue and PDO) tended to be mostly composed of more complex clones, both in terms of the number of SNVs/INDELs carried and the number of CNVs present in the dominant copy number clones in these samples (*i.e.* the tetraploid clone a in JK136, clones b, j, and m in JK142). Together, analyses of the bWES and scWGS data indicated that PDOs retained the genomic heterogeneity observed in their parent tumors, as evidenced by the comparable mutation- and copy number-based clonal frequencies found in tissue/PDO pairs.

### PDOs but not BTIC lines retain some non-malignant cells present in their parent tumors

To determine the extent to which PDOs retained phenotypic heterogeneity in addition to genetic heterogeneity, we next performed single-cell RNA sequencing (scRNA-seq) on tissue sections (n = 20, 1-2 sections per region) and PDOs (n = 57, 1-3 PDOs per region) from 10 patients (Supplementary Table 1). To further compare different model systems, we additionally performed scRNA-seq on BTIC lines (n = 9, one line per region) from a subset of five patients. We analysed 29,543 tissue cells, 35,395 PDO cells, and 10,137 BTICs that passed quality control filtering (Methods), with comparable medians of 2,526, 2,809, and 2,678 genes detected per cell, respectively. Visualizations of these cells using uniform manifold approximation and projection (UMAP) (McInnes et al., 2018) revealed similar patterns across sample types, most strikingly that cells from individual patients appeared to be more similar to each other than to cells from other patients (Figure 3A-C). This result is consistent with recent studies showing that inter-patient differences dominate clustering results at the single-cell level in GBM (Couturier, 2018; Darmanis et al., 2017; Jacob et al., 2020; Patel et al., 2014). However, we also observed cell groupings populated by cells from multiple patients in tissues and PDOs (Figure 3A-B, left, circled groups). These groupings displayed higher expression levels of non-malignant cell markers, including the immune cell marker *CD45*, the cancer-associated fibroblast (CAF) markers *ACTA2* and *PDGFRB* (Clavreul et al., 2014; Trylcova et al., 2015), the endothelial cell marker *VWF* (Soda et al., 2011), and the mature oligodendrocyte markers *MOG* and *MAG* (Figure 3A-B). These markers were either not detected or did not display distinct patterns of expression indicative of populations of non-malignant cells in BTICs (Supplementary Figure 3), consistent with the notion that there were no non-malignant cells retained in the BTIC lines, or at least none of suitable abundance for profiling.

**Figure 3:**
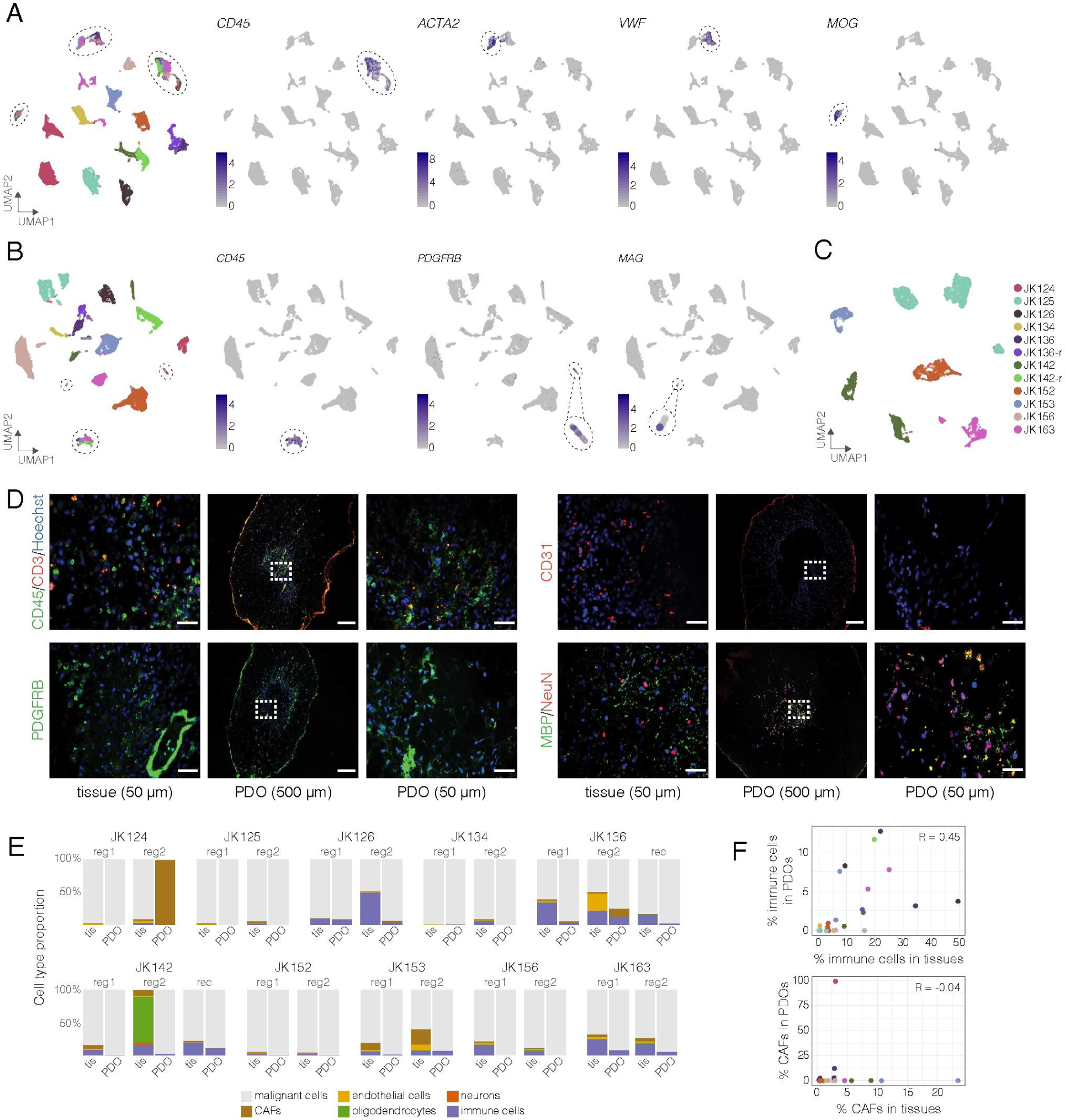
PDOs retain non-malignant cells. **A-C** UMAP visualization of all tissue cells (**A**), PDO cells (**B**), and BTICs (**C**), colored by patient (legend shown in panel C). For tissue and PDO cells, groupings of cells from multiple patients are circled, and expression of non-malignant cell markers is shown in these groupings on the right. *CD45*: immune cells; *ACTA2*/*PDGFRB*: CAFs; *VWF*: endothelial cells; *MOG*/*MAG*: mature oligodendrocytes. **D** Representative IF images showing the presence of immune cells (CD45; JK163 reg1), CAFs (PDGFRB; JK153 reg2), endothelial cells (CD31; JK153 reg1), and mature oligodendrocytes (MBP; JK136-r) in PDOs. White boxes on PDOs indicate the area shown in the zoomed-in images to their right. Hoescht staining is shown in blue in all images. Scale bar lengths are indicated below the images. **E** Proportion of tissue and PDO cells from each tumor region assigned to the cell types indicated. **F** Proportion of immune cells and CAFs in matched tissues and PDOs (each point represents a tumor region). Pearson correlation values are indicated. Color legend as in panel C.

To further study these populations of apparently non-malignant cells, we pooled data from each sample type independently and performed clustering analyses, identifying 27 tissue clusters, 33 PDO clusters, and 20 BTIC clusters (Supplementary Figure 3). In concordance with the UMAP visualizations, clusters were composed largely of cells from individual patients in all sample types, once again confirming the extensive level of inter-patient heterogeneity characteristic of GBM. In tissues and PDOs, clusters composed of cells from multiple patients were hierarchically distinct from other clusters and exhibited similarity to macrophages (tissue clusters t19, t4, and t20; PDO clusters o21 and o27), CAFs (t12 and o29), and endothelial cells (t14) as revealed by a similarity analysis to reference cell types (Methods). An examination of genes whose high expression characterizes these clusters (*i.e.* marker genes) included the markers shown in Figure 3A-B and further supported our results: for example, the immune cell clusters were marked by high expression of *e.g. AIF1*, *CD74*, *MNDA* and *CSF1R*; the CAF clusters displayed high expression of *DCN*, *BGN*, *IGFBP7*, and collagen genes; *PECAM1* (*CD31*), *ENG*, and *ECSCR* were highly expressed in cells from the endothelial cluster; and the oligodendrocyte cluster (t17) was marked by high expression of *MBP*, *MOBP*, *MOG*, and *MAG* (Supplementary Tables 4-6). To confirm that these non-malignant cell types were in fact observed in PDOs, we also performed immunofluorescence (IF) imaging and found evidence of the presence of immune cells (CD45), CAFs (PDGFRB) and mature oligodendrocytes (MBP; Figure 3D). Interestingly, while endothelial cells were only observed in tissues and not in PDOs in the scRNA-seq data, CD31 staining showed the presence of endothelial cells in PDOs as well, as previously reported (Jacob et al., 2020). Given their apparently low prevalence, it is possible that these cells were not of sufficient abundance to be reliably sampled by scRNA-seq.

To confirm the distinction between malignant and non-malignant cells using an orthologous approach, we also inferred copy number profiles from the scRNA-seq data (Methods). We first performed this analysis on the tissue cells using bulk RNA-seq data as a reference, and found that the candidate non-malignant cells clustered separately from the candidate malignant cells and that the latter displayed large-scale copy number variants characteristic of GBM such as amplification of chromosome 7 and deletion of chromosome 10 (Supplementary Figure 4). Notably, all cells from cluster t27 (35 cells) also clustered with non-malignant cells. Further examination of this cluster’s marker genes indicated that these cells were likely neurons, marked by expression of *e.g. NRCAM*, *NCAM2*, and *SHISA9* (Supplementary Table 4). Cells that were consistently identified as malignant or non-malignant by expression-based clustering and by inferred copy number clustering were retained for downstream analyses (29,499 of 29,543 cells, 99.9%). The analysis was then repeated for PDO and BTIC cells using the confirmed non-malignant tissue cells as a reference. In PDOs, a similar pattern was observed where candidate non-malignant cells clustered together and lacked characteristic GBM alterations that were found in candidate malignant cells. As with the tissue cells, the vast majority (35,382 of 35,395 cells, 99.96%) of cells were consistently identified as malignant or non-malignant and were retained for downstream analyses. Conversely, BTICs all appeared to carry copy number alterations, once again confirming the lack of non-malignant cells profiled in these samples.

Overall, we found that some tumors exhibited appreciable differences in the levels and composition of non-malignant cells across spatially distinct tissue samples from the same patient (Figure 3E). Although this variability could reflect sampling differences rather than differences in stromal levels across these tumors, replicate samples tended to display similar cell type proportions (Supplementary Figure 4), indicating that these were reproducible across samples. One striking example was found in the JK142 reg2 tissue section, which was composed mostly of non-malignant oligodendrocytes across replicate samples. Visual inspection of the original tumor section corroborated this observation, as it appeared to be largely composed of white matter (Supplementary Figures 4 and 5). This was also consistent with bWES and scWGS results that indicated low malignant cell proportions in independent samples from this region (0.28% and 1.6%, respectively). IF imaging of additional JK142 tissue samples revealed variable levels of MBP+ cells (Supplementary Figure 5; *e.g.* the reg2 sample has regions of high and low MBP staining), highlighting the extent of cellular heterogeneity that can be observed even within a relatively small region of a single tumor. Samples from JK124 reg2 showed an inverse relationship, where the tissue samples were largely composed of malignant cells, but the profiled PDO sample was almost entirely (118/119 cells) composed of CAFs. Notably, cell recovery from this particular PDO sample was poor, as reflected in the low number of cells profiled; therefore, it is possible that this result reflects a particular enrichment of CAFs during cell recovery from this sample rather than an accurate representation of the full spectrum of cells that were present in the PDO. Consistent with this notion, H&E staining of the JK124 reg2 PDO did not reveal abundant fibroblast-like morphology, and IF imaging indicated only moderately higher proportions of PDGFRB+ cells in the reg2 PDO compared to the reg1 PDO (Supplementary Figure 5). Nevertheless, bWES profiling of independent PDO samples from JK124 reg2 also indicated a low proportion (21%) of malignant cells. Apart from these two examples, PDOs generally appeared to harbor similar cell type proportions as their parent tumors, although the proportions of non-malignant cells were consistently lower in PDOs. We found that the proportions of immune cells in matched tissue/PDO pairs (Figure 3F) were correlated, indicating that PDOs derived from tissues with larger immune cell proportions were themselves likely to harbor more immune cells. Conversely, the relative number of CAFs was consistently low in PDOs (except for JK124 reg2 as discussed above) and was not correlated with the proportion of CAFs found in their parent tumors.

**Figure 4:**
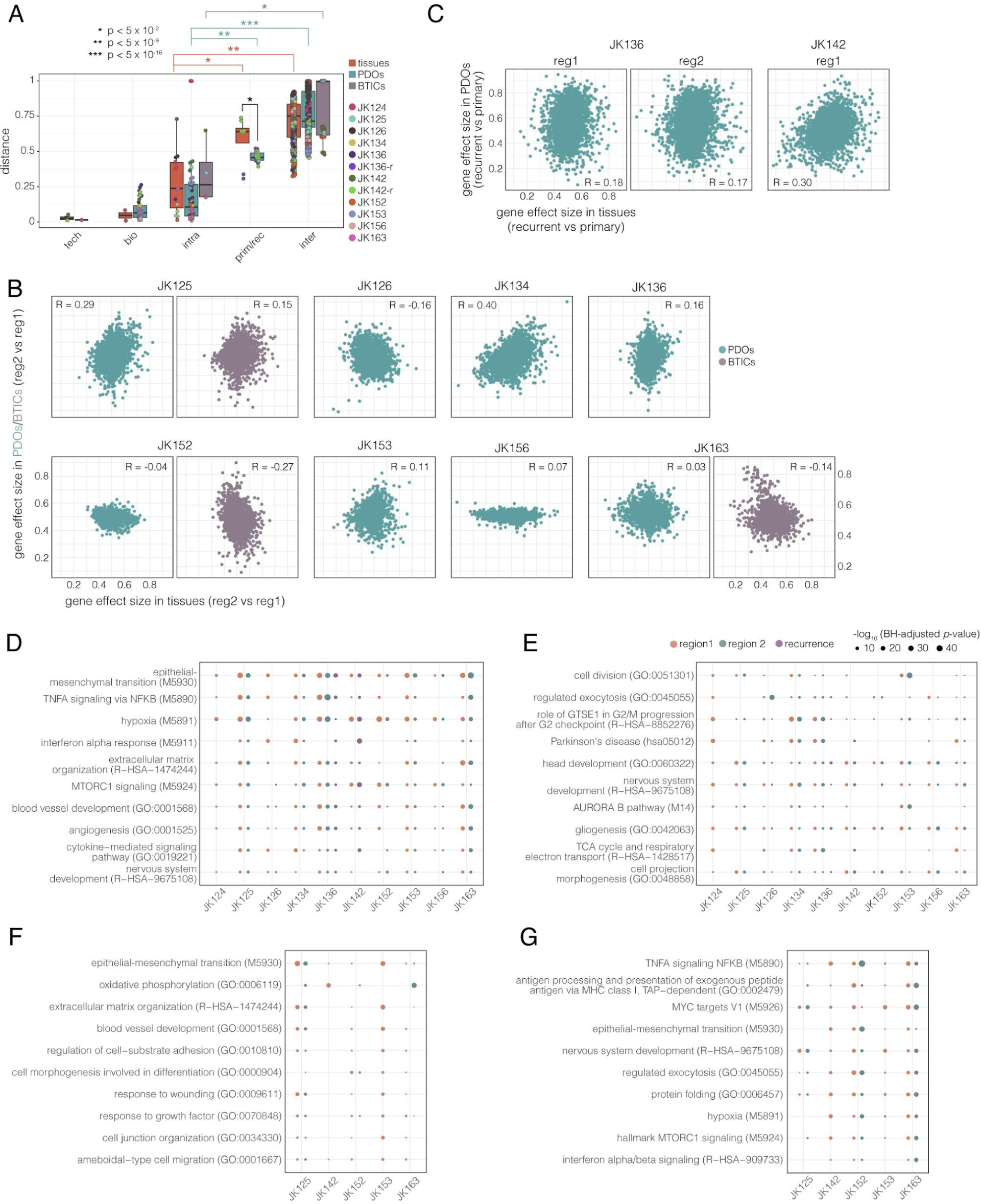
PDOs retain patterns of intra- and inter-tumor heterogeneity. **A** scUniFrac distances between different replicate types across tissue, PDO, and BTIC samples. Tech: same cell suspension run on different channels of the same 10X chip; bio: distinct tissue pieces or PDOs from the same tumor region; intra: samples from different regions of the same tumor; prim/rec: samples from matched primary and recurrent tumors; inter: samples from different patients. Indicated p-values were obtained using Wilcoxon tests. **B** Effect sizes of all detected genes in reg2 vs reg1 tissue cells (x-axis) and PDO or BTIC cells (y-axis). Effect sizes were calculated as overlap proportions from Wilcoxon rank sum tests (Methods), and only malignant cells were used for the analysis. Pearson correlation values are indicated for each comparison. **C** Effect sizes as in panel B in primary vs recurrent tissue cells (x-axis) and PDO cells (y-axis). **D-G** Top ten terms enriched for genes over-expressed (**D**) or under-expressed (**E**) in PDOs compared to tissues, or over-expressed (**F**) or under-expressed (**G**) in BTIC lines compared to tissues (Methods). Dots reflect the BH-adjusted *p*-value of each term across regional pairs. JK124 reg2 and JK142 reg2 were excluded since only one malignant cell was found in the PDO and tissue samples, respectively.

**Figure 5:**
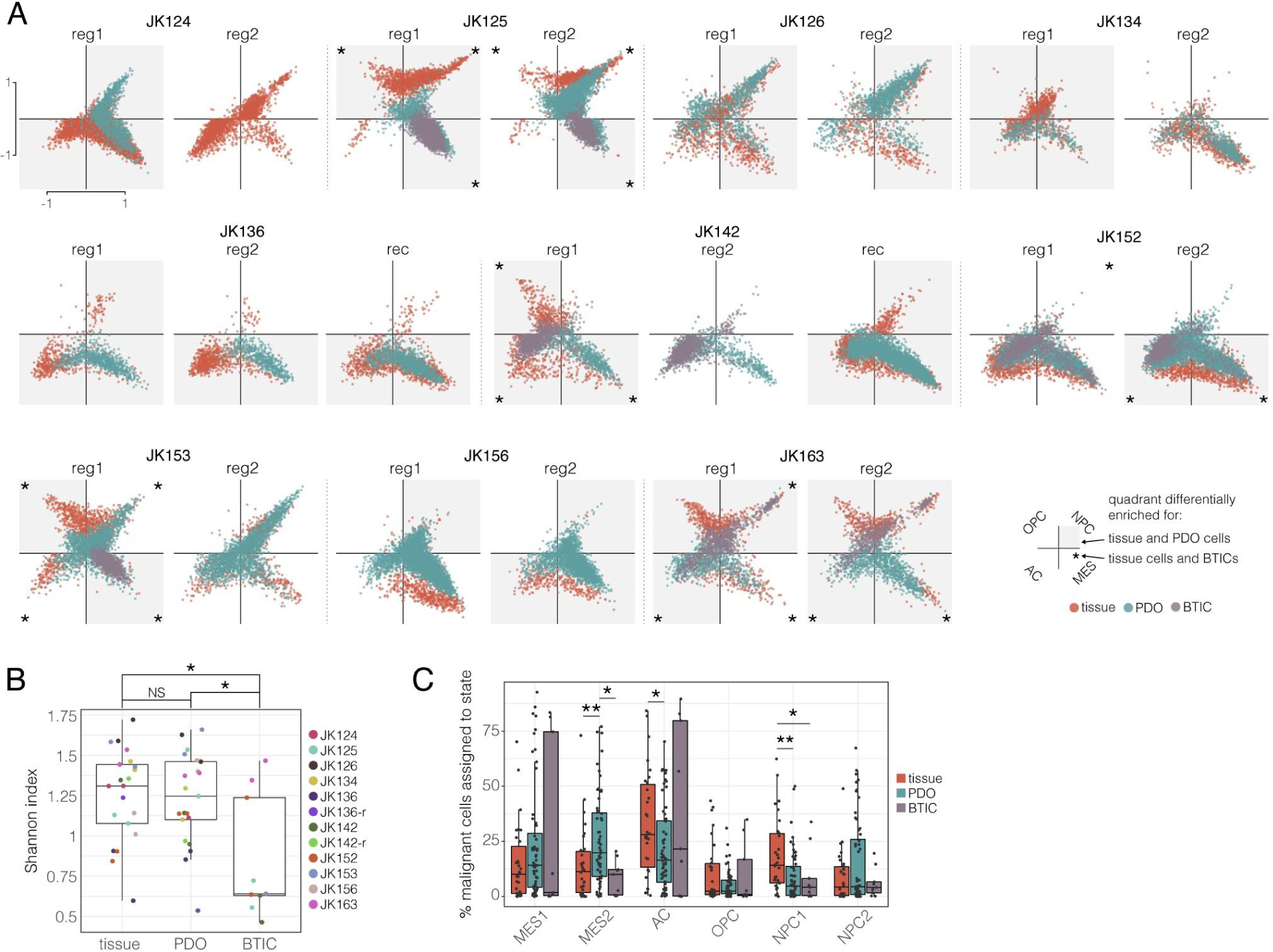
PDOs largely preserve cell state classifications relative to their parent tumors but have distinct distributions. **A** Two-dimensional representation of cellular states for all malignant cells (dots) from each sample type. Quadrants that are differentially enriched for tissue and PDO or tissue and BTIC cells are indicated by grey squares and asterisks, respectively (Fisher’s exact test *p* < 0.05). **B** Shannon entropy index of cell state assignment of malignant cells from regional samples. **C** Proportion of cells from each sample assigned to each cellular state. \**p* < 0.05, \*\**p* < 0.01 (Wilcoxon rank sum test).

### PDO macrophages have distinct profiles compared to those in tumors

To explore the composition of immune cells found in the PDOs and their relationship to those observed in matched tissue samples, we first used UMAP to visualize the combined tissue and PDO immune cells (3,224 tissue and 1,042 PDO immune cells; Supplementary Figure 6). The PDO immune cells largely clustered apart from the tissue immune cells, except for one small group, composed of 22 tissue cells and 10 PDO cells, that expressed T-cell markers such as *CD3D/E/G* (group 1; Supplementary Table 7). This is consistent with an observation of T-cells present in both tissue and PDO samples as shown by IF (Figure 3D). *CD8A/B* and cytotoxic markers such as *GZMA/B/H* and *IFNG* were also highly expressed in this group, indicating that it was likely predominantly composed of cytotoxic CD8 T-cells. Conversely, cells in groups 2 through 5 showed relatively high expression of markers of glioma-associated macrophages/microglia (GAMMs) (Muller et al., 2017). We also confirmed the presence of AIF1/IBA1+ and CD68+ GAMMs in the PDOs using IF (example shown in Supplementary Figure 6). The fact that T-cells from tissues and PDOs cluster together while GAMMs from the two sample types cluster separately is consistent with the notion that the latter observation is likely due to biological differences rather than technical effects, as these would presumably be observed across cell types if they were present.

A differential expression analysis between the four GAMM groups revealed that cells in group 2 were characterized by high expression of genes involved in oxidative phosphorylation (*e.g. ATP5PB*/*PO*/*MC1*/*MC3*/*F1A*/*F1C*) and cell cycle (*e.g. CDK1* and *EIF4EBP1*); group 3 cells were marked by high expression of multiple complement genes (*e.g. C1QA*/*B*/*C*, *C3*); cells in group 4 were marked by high expression of genes involved in extracellular matrix (ECM)-modulating factors, including for example matrix metallopeptidases (*MMP1*/*9*/*12*/*14*), C-type lectin domain family genes (*CLEC4A*/*4E*/*7A*) and several cytokines (*CXCL1/3/5/8*, *CCL2/7/8*, *IL1B*, *IL1RN*); and group 5 cells had high expression of genes activated in response to tumor necrosis alpha (TNFɑ) signaling (*e.g. TRAF1* and *BIRC2*) as well as *PD-L1* (*CD274*). Interestingly, *PD-L1* was detected in ∼10% of PDO immune cells (107/1,042) but only in <1% of tissue immune cells (12/3,224). Together, these results indicated that the immune cells retained in the PDOs included T-cells and macrophages that interact with their environment and express many of the markers found in tissue immune cells, which is consistent with the notion that these cells are active and functional.

### PDOs retain inter- and intra-tumor patterns of heterogeneity

The UMAP visualizations and clustering results from tissues and PDOs revealed a similar observation in both contexts: while stromal cells from different patients were sufficiently similar to each other to cluster together, malignant cells displayed more pronounced inter-patient transcriptional differences. To quantify these observations, we used scUniFrac (Liu et al., 2018) to calculate distances between all pairs of samples based on the hierarchical clustering results in each sample type. As expected, distances between inter-patient tissue pairs were significantly larger than distances between intra-patient (*i.e.* reg1 and reg2) tissue pairs; notably, similar patterns were also observed in both PDOs and BTICs, indicating that both model systems retain intra-tumor similarities and inter-tumor differences (Figure 4A). Distances between primary/recurrent pairs in both tissues and PDOs were larger than those between intra-patient pairs but smaller than those between inter-patient pairs, indicating that recurrent tumors retained transcriptional features reminiscent of their matched primary tumors, and that PDOs derived from these tumors reflected these observations. Notably, distances between matched tissue samples that were processed in independent experiments were similar to those between samples that were processed in the same experiment and on the same 10X chip, indicating minimal between-chip batch effects (Supplementary Figure 7). Conversely, PDOs did show higher distances between cross-chip pairs; however, PDO pairs were only available for two patients—namely JK125 and JK136—and the cross-chip sample for JK136 showed a starkly different cell type composition compared to the other JK136 PDOs (reg2 PDO #4: 25.9% malignant cells vs a mean of 94.1% malignant cells in the other JK136 PDOs; Supplementary Figure 4). Given that non-malignant cells cluster separately from malignant cells, this difference can explain the larger distances seen in the JK136 cross-chip comparison; therefore, the observation of higher cross-chip distances between PDOs compared to tissues may be due to the specific samples examined rather than to a technical effect at play specifically in the PDOs.

To better understand patterns of intra-tumor heterogeneity and how these are recapitulated in PDOs and BTICs, we characterized spatial transcriptional heterogeneity. To do this, we performed differential expression (DE) analyses between malignant cells from different regions for each patient, independently for each sample type (*i.e.* JK125 tissue cells reg1 vs reg2, JK125 PDO cells reg1 vs reg2, JK125 BTICs reg1 vs reg2, and so on; Methods), and then compared results between sample types. We observed considerable variability between tumors: for some patients (*e.g.* JK125 and JK134), DE patterns between regional PDOs tended to replicate DE patterns between regional tissue samples (*i.e.* genes that were more highly expressed in reg2 tissue cells compared to reg1 tissue cells were also more highly expressed in reg2 PDO cells compared to reg1 PDO cells, and vice-versa), while for other patients we did not observe consistent patterns between pairs of tissues and pairs of PDOs (Figure 4B). This is consistent with the notion that malignant cells within PDOs can retain intra-tumor expression patterns found in the parent tumors, but do not necessarily do so consistently. Interestingly, in cases where both PDOs and BTICs were available, the effect size correlations were consistently lower between tissues and BTICs than between tissues and PDOs; therefore, despite the variability observed in PDOs’ ability to retain intra-tumor transcriptional differences, these appear to be consistently better represented in PDOs than in BTICs. Notably, comparisons of primary/recurrent pairs yielded similar results; specifically, JK136 primary/recurrent pairs had correlations comparable to JK136 regional pairs, and the JK142 primary/recurrent comparison had the highest correlations observed across all pairs examined (Figure 4C), indicating that PDOs were also able to retain some recurrence-specific gene expression patterns.

### Both model systems are subject to systematic effects on gene expression

To explore whether the culture conditions used to propagate PDOs and BTICs induced consistent changes in gene expression, we first performed DE analyses comparing malignant cells from each regional tissue/PDO and tissue/BTIC pair, then performed enrichment analyses with an emphasis on identifying commonly enriched terms (Methods). Some terms were commonly identified across regional pairs: for example, genes more highly expressed in PDOs compared to tissues were commonly enriched for processes such as the epithelial-to-mesenchymal transition (EMT), hypoxia, and ECM organization, while genes with lower expression in PDOs compared to tissues were enriched for cell cycle processes and oxidative phosphorylation (Figure 4D-E and Supplementary Tables 8-10). Interestingly, genes that were more highly expressed in PDOs compared to tissues were also enriched for cytokine responses, including TNFɑ and interferon signaling (*e.g. B2M*, *BCL6*, *CD44*, *HLA-A/B/C*, *IFI27/35*, *IFIT1/3*, *IRF1/7*, and *TNFAIP8*), indicating the possibility that such cytokines are generally present in higher levels in PDOs compared to the sampled tissues. Cytokine genes and genes activated in response to TNFɑ signaling were also more highly expressed in PDO GAMMs compared to tissue GAMMs (discussed above, Supplementary Figure 6), possibly indicating global PDO-related effects or interactions that amplify these responses between malignant cells and GAMMs within PDOs.

Compared to PDOs, BTICs showed less consistent gene expression changes relative to the parent tissues. For example, both over-expressed and under-expressed genes in BTIC lines compared to tissues were enriched for EMT (Figure 4F-G). Some genes involved in EMT were in fact over-expressed in some BTIC lines compared to their parent tissues and under-expressed in other lines: for example, *CD44* and *VIM* were significantly over-expressed in the JK125, JK153, and JK163reg2 lines, but significantly under-expressed in the JK142 and JK152 lines (FDR < 0.05; Supplementary Table 9). This indicates that, at least for EMT, which plays an important role in cancer progression (Dongre and Weinberg, 2019), BTIC lines do not appear to have consistent differences in gene expression compared to their parent tissues. Genes with lower expression in BTICs compared to tissues consistently yielded enrichment for processes dependent on or involved in malignant cell interactions with non-malignant cells, including cytokine signaling (*e.g.* TNFɑ and interferon signaling) and antigen processing and presentation. This under-expression may reflect the lack of non-malignant cells and therefore lack of malignant-to-non-malignant cell interactions within BTIC lines. Together, these results indicate that many of the gene expression changes between tissues and either PDOs or BTICs are related to cell-extrinsic interactions and thus may largely reflect the distinct microenvironments found in each sample type; however, these differences appear to be more consistent in PDOs compared to BTIC lines.

### PDOs largely recapitulate cell state distributions observed in parent tissues

To further explore and characterize transcriptional heterogeneity within the malignant cell populations present in different sample types, we scored individual cells for recently described GBM-specific cell states (Neftel et al., 2019) and compared the distribution of these states between regions and sample types (Methods). Overall, the relative proportions of cells assigned to meta-states differed significantly (Fisher’s exact test *p* < 0.05) between matched tissues and PDOs or matched tissues and BTICs in most cases (Figure 5A): of 80 tissue/PDO meta-state pairs, 43 were differentially enriched (54%, gray squares), and of 32 tissue/BTIC meta-state pairs, 21 were differentially enriched (66%, asterisks). While PDO cells appeared to retain heterogeneous profiles relatively similar to cells in their parent tissues despite their recurrent differential enrichment, BTICs appeared to largely coalesce into one or two meta-states (*e.g.* JK125 BTICs were almost entirely assigned to the mesenchymal meta-state; Supplementary Figure 7). To quantify this observation, we calculated entropy values for each sample type in each tumor region based on the distribution of cells assigned to each cell state and indeed found that PDO cells had similar entropy as the parent tissues, while BTICs had significantly (Wilcoxon rank sum test *p* < 0.05) lower entropy than both tissue and PDO cells (Figure 5B). Together, these results indicate that, while PDOs do not have identical cell state distributions as their parent tumors, they are capable of displaying the same states as their parent tumors. Conversely, BTICs do not exhibit this capacity, with cells mostly converging onto the transcriptional profiles of one or two cell states.

Given the observations described above, we sought to investigate whether PDOs and BTICs exhibited systematic differences in the distributions of these GBM-specific cell states. To do this, we compared the proportions of cells assigned to each state within each sample. Compared to tissues, we found that PDOs tended to have higher proportions of cells assigned to the MES2 state and lower proportions of cells assigned to the AC and NPC1 states (Wilcoxon rank sum test *p* < 0.05; Figure 5C). Given that the MES2 state is hypoxia-dependent, the higher proportions of cells assigned to this state in PDOs is likely related to the overall increase in expression of hypoxia-related genes in PDO cells compared to tissue cells (Figure 4D). Hypoxic conditions are common in GBM and have been shown to contribute to these tumors’ aggressive nature (Colwell et al., 2017); however, hypoxia-induced genes tend to be most highly expressed in cells from tumor cores compared to cells from near the invasive edge (Darmanis et al., 2017). Given that the tissue cells used in this study were sampled from the enhancing edge and that the PDOs appear to form “miniaturized tumors” with their own cores (Figure 1), it is possible that the hypoxia signal we observe is a result of the increased likelihood of sampling hypoxic cells in the PDOs.

Interestingly, compared to tissues, both PDOs and BTICs had lower proportions of cells assigned to the NPC1 state (Wilcoxon test *p* < 0.05), which is defined by OPC lineage genes. Conversely, proportions of cells assigned to the NPC2 state, which is defined by neuronal lineage genes, were not statistically significantly different in either PDOs or BTICs, although the former did show a trend towards increased proportions compared to tissues. While this may indicate that the neuronal stem cell media used to propagate BTICs and PDOs biases neuronal progenitor-like cells away from an OPC-like profile, we note that the proportions of actual OPC-like cells do not differ significantly between tissues, PDOs, and BTICs. Therefore, if the culture conditions are indeed inducing this change, they may affect earlier progenitor-like cells. PDOs also contain lower proportions of astrocytic-like cells than tissues (Wilcoxon test *p* < 0.05), which may reflect a bias away from more differentiated cell types in this model.

This analysis also revealed unique bimodal distributions of cell proportions assigned to the MES1 and AC states in BTIC lines, whereas these states showed a more continuous distribution in tissues and PDOs. Indeed, BTIC lines appeared to be composed almost entirely of mesenchymal cells (JK125 and JK153), astrocytic cells (JK142 and JK152), or a mix of NPC- and OPC-like cells (JK163; Supplementary Figure 7). Interestingly, this did not always correlate with the major cell states observed in the parent tissues: for example, tissue cells from both regions of JK125 were largely assigned to NPC- and OPC-like states (97.4% [1,539/1,580 cells] in reg1 and 96.2% [1,129/1,173] in reg2), whereas BTICs from these regions were, as mentioned previously, almost entirely assigned to one of the mesenchymal states (95.7% [2,464/2,575] in reg1 and 93.2% [1,479/1,587] in reg2). Neftel *et al*. (2019) also showed that a proneural-to-mesenchymal transition in BTICs was associated with recurrence and shorter survival. Tissue samples from JK125 and JK153 both had strong proneural signatures (96.9% and 66.5% of malignant cells assigned to an OPC- or NPC-like state, respectively) whereas the BTIC lines derived from these tumors both had strong mesenchymal signatures (94.7% and 93.3% of BTICs assigned to a mesenchymal-like state, respectively). While proneural signatures in tumors are typically associated with relatively favorable outcomes, including a trend towards longer survival, compared to tumors with other transcriptional signatures (Verhaak et al., 2010; Wang et al., 2017), these two patients had the shortest survival within our cohort (69 days and 194 days post-diagnosis, respectively).

## Discussion

Representative cell-based models are crucial to studying cancers and properly understanding their underlying biology. While neurosphere cell lines have been used to study GBM for more than two decades and have provided useful insights, the emergence of more complex models, including xenografts and organoids, have begun to allow for the dissection of GBM heterogeneity *in vitro*. However, the cellular composition of such models needs to be comprehensively mapped in order to understand the nature of the research questions that can be addressed using them. Here, we have used bulk and single-cell genomic approaches to perform an in-depth analysis of the genetic and transcriptional heterogeneity observed in GBM, novel GBM PDOs, and BTIC lines. At the genetic level, matched tissues and PDOs were consistently at least as similar to each other than were tissue sections from opposite regions of the same tumor. Subclonal distributions, either defined at the mutational or copy number level, were also comparable between tissue/PDO pairs, indicating that PDOs accurately reflect the genomic profile of the tissue section from which they were derived.

In addition to the retention of genetic characteristics, PDOs also appeared to retain much of the transcriptional heterogeneity observed in the parent tissues. Firstly, the scRNA-seq analysis revealed that PDOs retained some non-malignant cell types, namely immune cells, CAFs, and oligodendrocytes. This raises the possibility that PDOs could provide useful models to study tumor/microenvironment interactions, which could also be enhanced by *e.g.* co-culturing with isolated peripheral immune cells from matched patients (Yuki et al., 2020). However, the transcriptional profiles of PDO macrophage populations differed substantially from those of tissue macrophage populations, and functional studies of these populations will be needed to further elucidate how well they represent *in vivo* GBM-associated immune cells.

Despite the retention of some non-malignant cell types, PDOs are predominantly composed of malignant cells. While these generally resemble the malignant cells observed in tissues, they do not appear to fully represent the gene expression programs at work in tissues. For example, PDO cells tended to consistently display higher expression of genes involved in hypoxia, EMT, and ECM organization. Some of these effects may be mediated by differences in the structures being sampled; for example, the increased expression of hypoxia genes may be due to the fact that entire PDOs were dissociated for profiling, and thus cells from the hypoxic core were likely to be sampled, whereas the tissue cells profiled in this study were obtained from peripheral tumor sections away from hypoxic cores and were thus less likely to be expressing markers of hypoxia response (Darmanis et al., 2017). Other effects, such as the increased expression of genes involved in ECM modulation, may reflect the influences of the environment in which the PDOs are grown. Interestingly, genes commonly expressed at lower levels in PDOs compared to tissues were enriched for cell cycle processes and oxidative phosphorylation, indicating the possibility that decoupling GBM cells from their native environment influences gene expression programs.

Recently, Neftel et al. (2019) used scRNA-seq approaches to show that transcriptional heterogeneity in glioma consistently converged onto six defined cellular states, in addition to cell cycle states. Distributions of PDO cells assigned to each of these states were similar to those observed in tissues, indicating that the PDO growth conditions do not consistently enrich for or deplete specific states. States that did show differential enrichment included MES2, which is a hypoxia-dependent mesenchymal state, as well as NPC1, which is defined by the expression of genes from the oligodendrocyte precursor lineage. The higher proportions of MES2 cells in PDOs is likely related to the increased sampling of hypoxic cells, as discussed above. Conversely, the lower proportions of NPC1 cells found in PDOs compared to matched tissues may be a consequence of culture media supplementation with EGF and FGF-2, which supports the growth of neural stem cells but has been shown to lead to the loss of OPC markers in culture (Ledur et al., 2016). Notably, a reduction in the proportion of NPC1-assigned cells compared to tissues was also seen in BTICs, as reported by Ledur et al. (2016). The proportions of cells assigned to the OPC state, however, did not differ significantly between tissues, PDOs, and BTICs, indicating that the culture conditions may affect more primitive cells, perhaps destined to become OPCs, rather than OPC-like cells themselves.

As part of this study, we were also able to generate and profile PDOs from two recurrent tumors. While the molecular profiles of primary GBMs have been extensively studied, mostly at the bulk level (*e.g.* through The Cancer Genome Atlas [TCGA] efforts (Brennan et al., 2013; Ceccarelli et al., 2016)) and to a lower extent through single-cell studies (Johnson et al., 2018), ultimately most patients succumb to recurrent disease. Given that recurrent tumors are often genetically and transcriptionally distinct from their primary tumors (Kraboth and Kalman, 2020), a better understanding of the molecular underpinnings of recurrent disease will likely be crucial to improving outcomes for GBM. Two main factors make the study of recurrent GBM tumors particularly difficult: (1) only ∼25% of patients are eligible for repeat surgery upon recurrence (Weller et al., 2013), and (2) even resectable recurrent tumors tend to have a higher rate of necrosis, which can make molecular profiling difficult or impossible (Marucci et al., 2015). Given that even a sample found to have low tumor content by multiple orthogonal methods (bWES, scWGS, and scRNA-seq) was able to seed successful organoids in this study (JK142 reg2), the use of PDOs may help address the second point in particular by creating useful models amenable for downstream profiling, even from small amounts of tumor material. Indeed, the organoids derived from the recurrent samples studied here appeared to faithfully represent their parent tumors as did the organoids derived from primary samples. For example, in both tissues and PDOs, cells from primary/recurrent pairs clustered together less often than cells from regional pairs, but more often than cells from different patients, indicating that the recurrent samples at least partially retained the transcriptional profiles of their matched primary samples. The data presented here thus also provide a rich resource to study recurrent tumors at single-cell resolution.

Overall, the PDOs profiled in this study appear to retain much of the phenotypic and genomic heterogeneity observed in GBM. In contrast to the commonly used BTIC models, they support the survival of non-malignant cell types and show a distribution of malignant cell states much more closely related to what is observed in tissue samples. Nevertheless, the PDOs do show some potentially important differences that need to be appreciated, and we showed that a single-cell profiling approach was key to dissecting this variability. Indeed, we have presented an integrative analysis of scWGS data, bWES data, and the largest single GBM scRNA-seq dataset available to date, which provided valuable insights into the similarities and differences between primary GBMs, PDOs, and BTIC lines. Our study has shed light on the biology of GBM and has helped highlight the strengths and weaknesses of PDO and BTIC models to address outstanding questions in the field.

## Acknowledgements

The authors gratefully acknowledge that this work would not have been possible without the participation of patients and their families. We thank the Calgary Brain Tumour and Tissue Bank for providing patient samples, and the Sequencing, Bioinformatics, and Project Management teams at Canada’s Michael Smith Genome Sciences Centre for expert technical contributions. We thank Dr Samuel Aparicio, Dr Andrew Roth, Ishika Luthra, and members of the Marra lab for helpful discussions. This study was supported by the Terry Fox Research Institute (TFRI; Translational Research Program Grant #2009-20) and the Canadian Institutes of Health Research (CIHR; FDN-143288). We are especially grateful to Ms Donna Anderson for her generosity in supporting this project. M.A.M. gratefully acknowledges the support of BC Cancer, the BC Cancer Foundation, the Canada Research Chairs program, the Canada Foundation for Innovation, Genome BC, Genome Canada, and the University of British Columbia (UBC). V.G.L. was supported by a CIHR Vanier Canada Graduate Scholarship, a Killam Doctoral Scholarship, and a UBC 4-year fellowship.

## Author Contributions

Conceptualization: M.A.M, J.J.K., and V.G.L.; Methodology: J.J.K, M.H., D.L., and V.G.L.; Formal Analysis: V.G.L.; Investigation: V.G.L., D.L.T., and S.A.; Resources: M.A.M., J.J.K., and J.C.; Data Curation: V.G.L.; Writing - Original Draft: V.G.L.; Writing - Review & Editing: V.G.L. and M.A.M.; Visualization: V.G.L. and S.A.; Supervision: M.A.M, J.J.K., and J.G.C.; Funding Acquisition: M.A.M, J.G.C., J.J.K., and M.B.

## Supplementary Figures

**Supplementary Figure 1:**
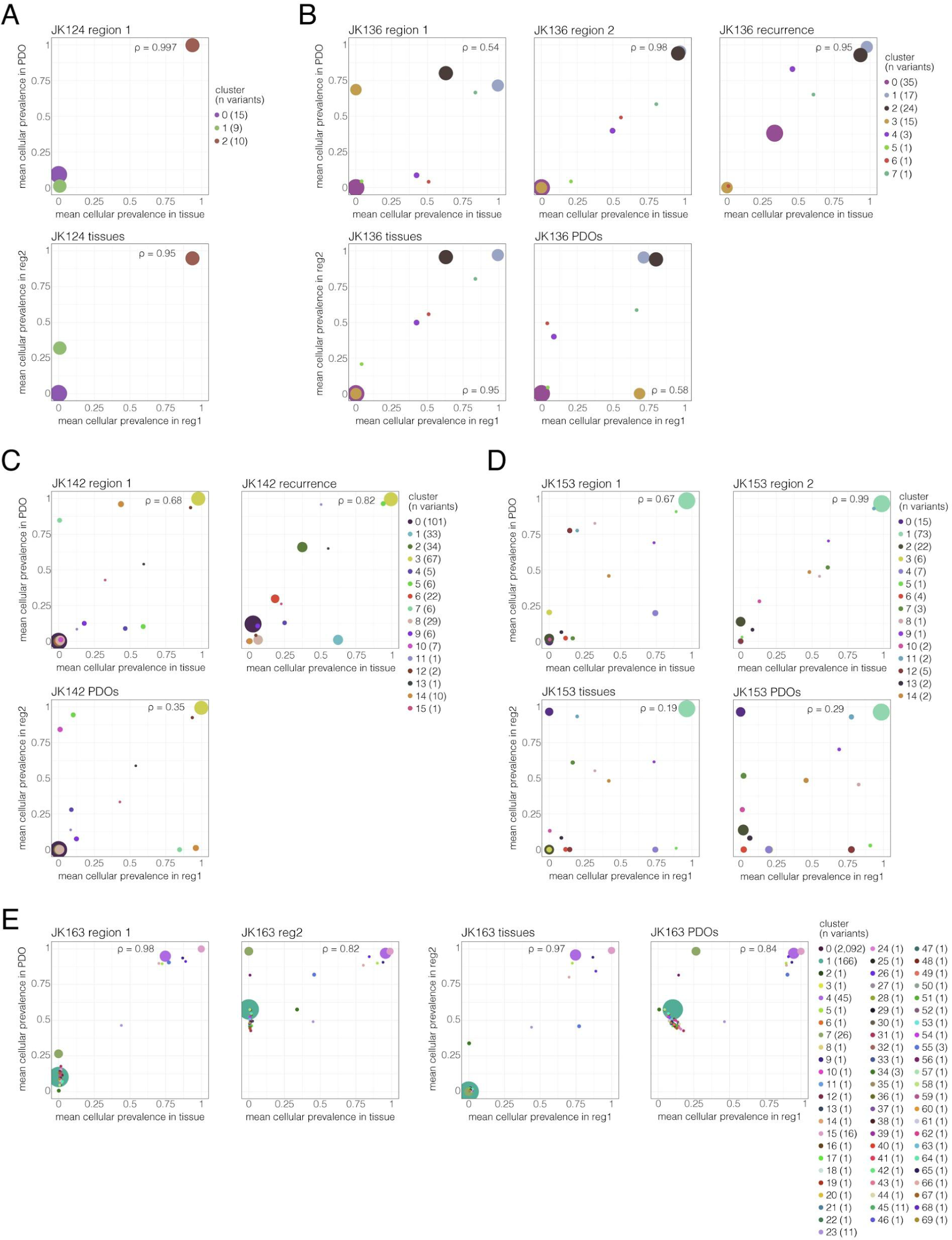
Correlations of variant clusters are comparable between tissue/PDO pairs and regional tissue or PDO pairs. Scatter plots showing cellular prevalence of variant clusters for tissue/PDO pairs (top for panels A-D, left for panel E) and regional tissue or PDO pairs (bottom for panels A-D, right for panel E), as obtained using PyClone (Roth et al., 2014). Dot area is proportional to the number of variants in each cluster, which is also indicated in the respective color legends. Spearman’s *rho* correlation values are shown on each plot.

**Supplementary Figure 2:**
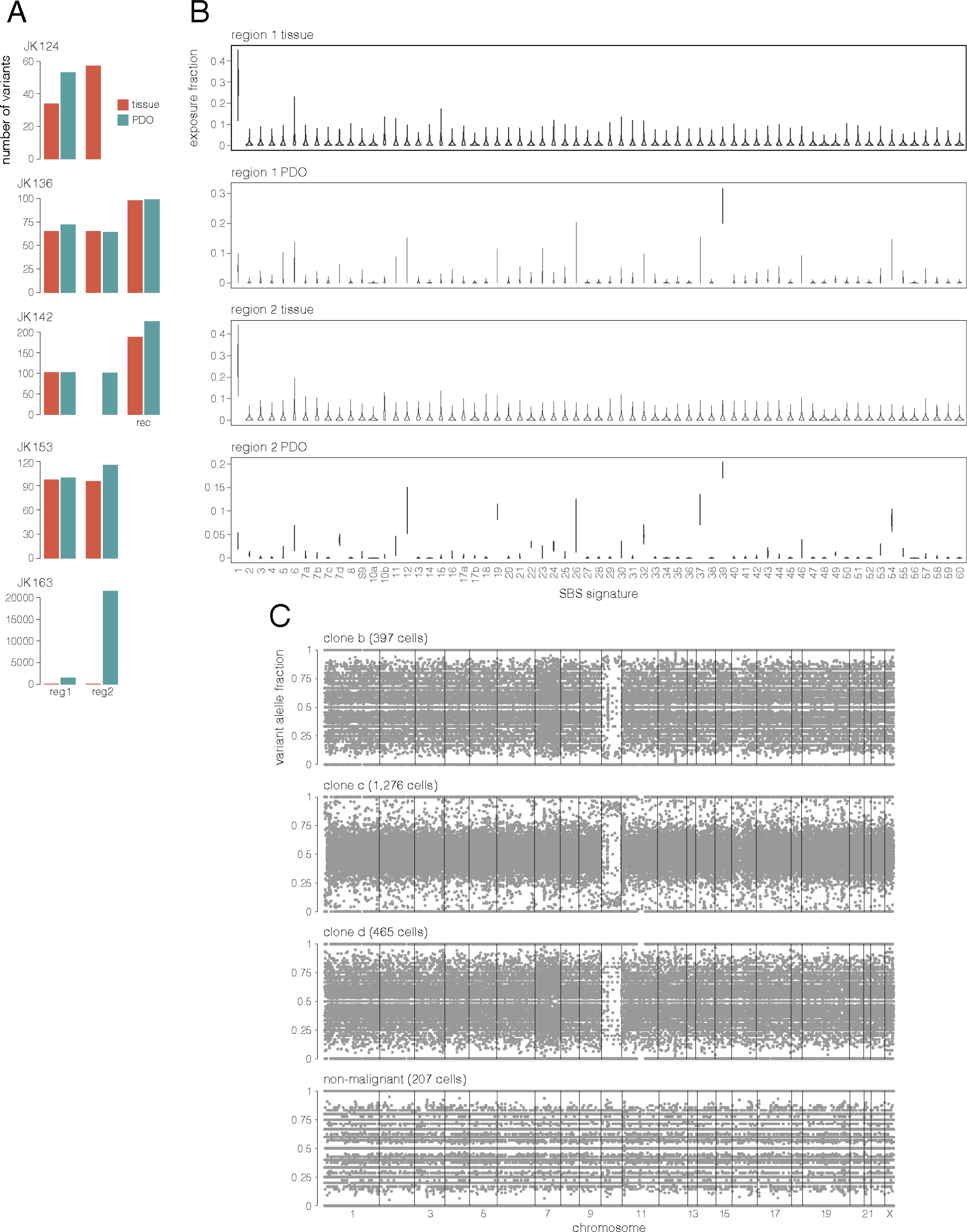
JK163 PDOs display mutation signatures not found in PDOs from other patients. **A** Number of variants identified using bWES in each sample (read depth ≥ 20 and alternate read depth ≥ 10). With the exception of JK163, PDOs appeared to carry a similar number of variants as their parent tumors. **B** Exposure fraction of variants identified using bWES across JK163 tissue and PDO samples against single base pair (SBS) signatures defined by the PCAWG consortium (Alexandrov et al., 2020), as calculated using SignIT (Zhao et al., 2017). SBS39 is enriched in both PDO samples, but not in the matched tissues. **C** Variant allele fractions of germline heterozygous mutations (identified using bWES) calculated from scWGS reads pooled across JK153 cells assigned to the same malignant clone or identified as non-malignant cells. LOH can be observed across chromosome 10 in all three malignant clones, indicating copy number neutral LOH in clone b cells.

**Supplementary Figure 3:**
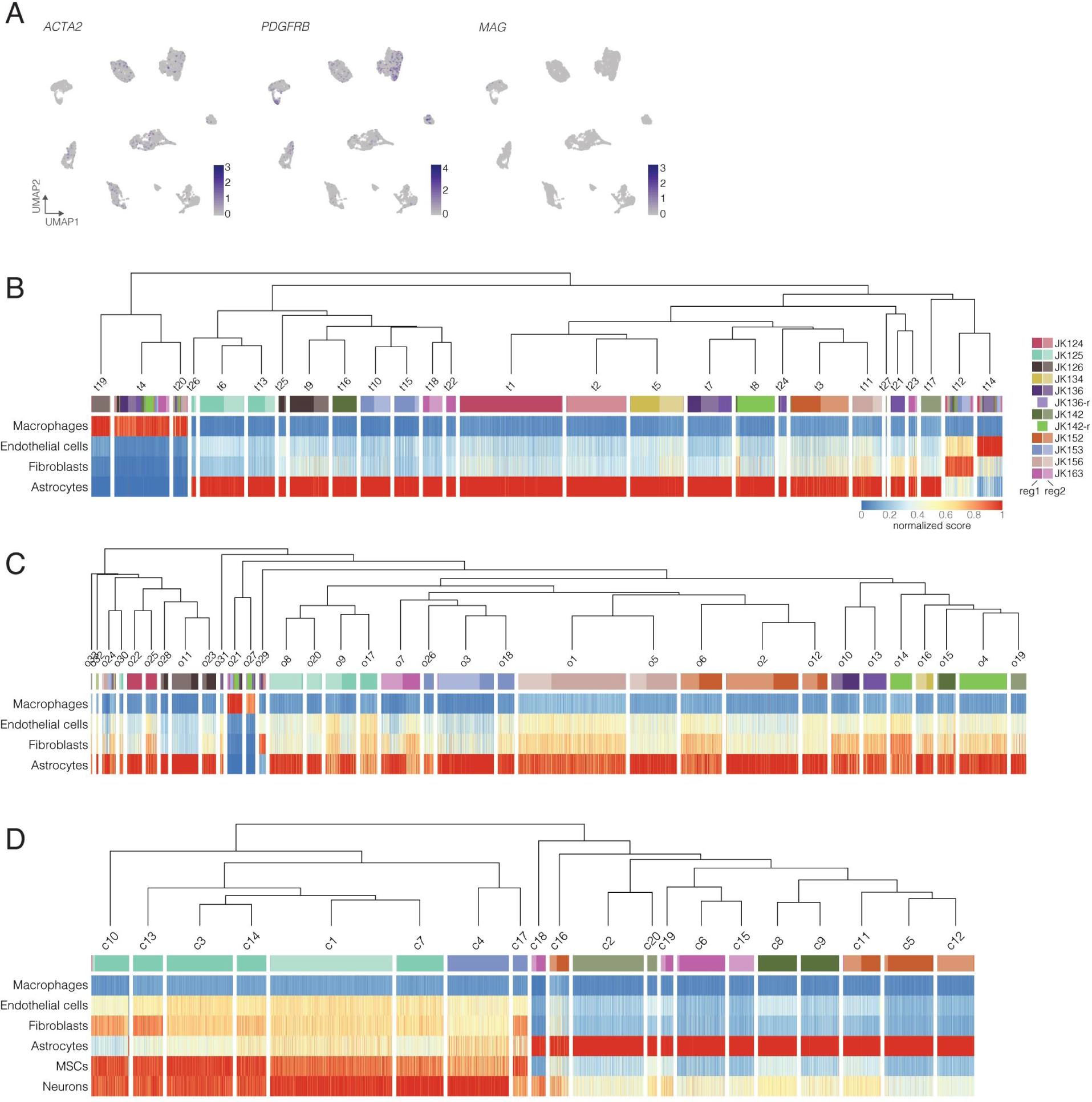
Non-malignant cells in tissues and PDOs cluster separately from malignant cells and have expression profiles similar to those of reference cell types. **A** UMAP visualizations of BTICs showing expression of non-malignant cell markers; populations of non-malignant cells are not evident as in tissues and PDOs. Other markers shown for tissue and PDO cells (*CD45*, *VWF*, and *MOG*) were not detected in this dataset. **B-D** Heatmap of normalized similarity scores for all tissue cells (B, 29,543 cells), PDO cells (C, 35,395 cells), and BTICs (D, 10,137 cells) compared to select reference cell types (Methods). Trees were obtained using hierarchical clustering of mean significant PC values across clusters (*i.e.* clustering of clusters, see Methods).

**Supplementary Figure 4:**
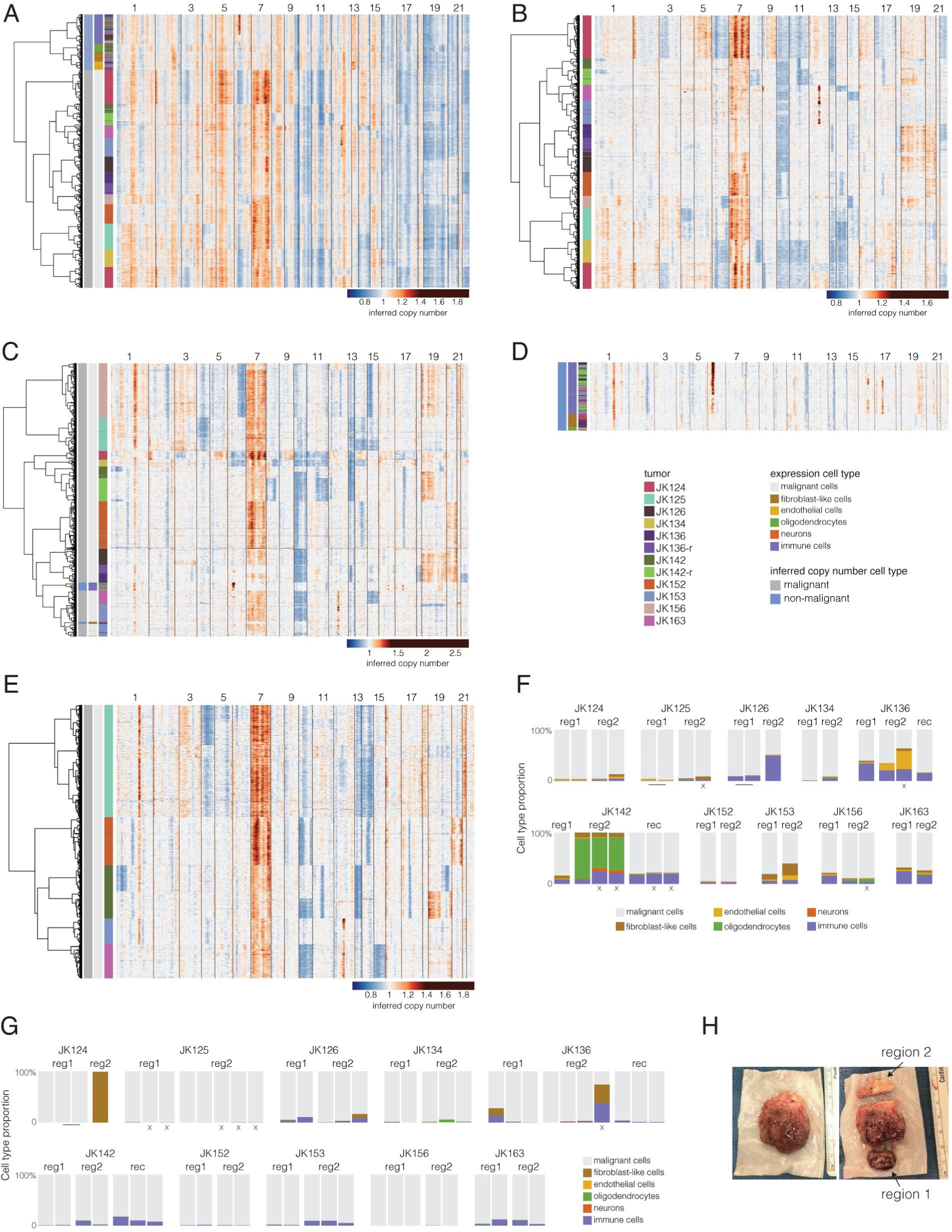
Copy number inference confirms the presence of non-malignant cells in tissues and PDOs, as well as their absence in BTICs. **A** Inferred copy number in all tissue cells obtained using infercnv and bulk profiles from GTEx normal brain samples as a reference (Methods). Candidate non-malignant cells (as identified by cell clustering shown in Supplementary Figure 3) cluster separately from candidate malignant cells and lack GBM-associated large copy number alterations (CNAs) that are observed in candidate malignant cells, such as amplification of chr7 or deletion of chr10. **B** Inferred copy number in all malignant tissue cells obtained using infercnv and non-malignant single cells as a reference. Using single cells as reference helps reduce noise and highlights large-scale CNAs observed in malignant cells (*e.g.* chr7 amplification, chr10 deletion). **C** Inferred copy number in all PDO cells, obtained by infercnv using non-malignant tissue cells as a reference. GBM-associated CNAs (*e.g.* chr7 amplification, chr10 deletion) are observed in candidate malignant cells. **D** Larger view of cells identified as non-malignant by clustering of inferred copy number (*i.e.* labeled as non-malignant in panel A) showing that GBM-associated CNAs are not observed in these cells. **E** Inferred copy number in all BTICs, obtained by infercnv using non-malignant tissue cells as a reference. All cells show typical GBM-associated CNAs, confirming the absence of non-malignant cells. **F-G** Proportion of cells from each tissue (F) and PDO (G) sample assigned to the indicated cell types by both expression and inferred copy number analysis. Technical replicates (*i.e.* two samples obtained from the same cell suspension) are linked by bars. Biological replicates run in independent experiments (*i.e.* cross-chip replicates) are indicated by an “x”. **H** Image of resected tumor from JK142. Region 2 appears mostly composed of white matter, which agrees with the scRNA-seq results indicating that the majority of cells from this sample were non-malignant oligodendrocytes (panel F).

**Supplementary Figure 5:**
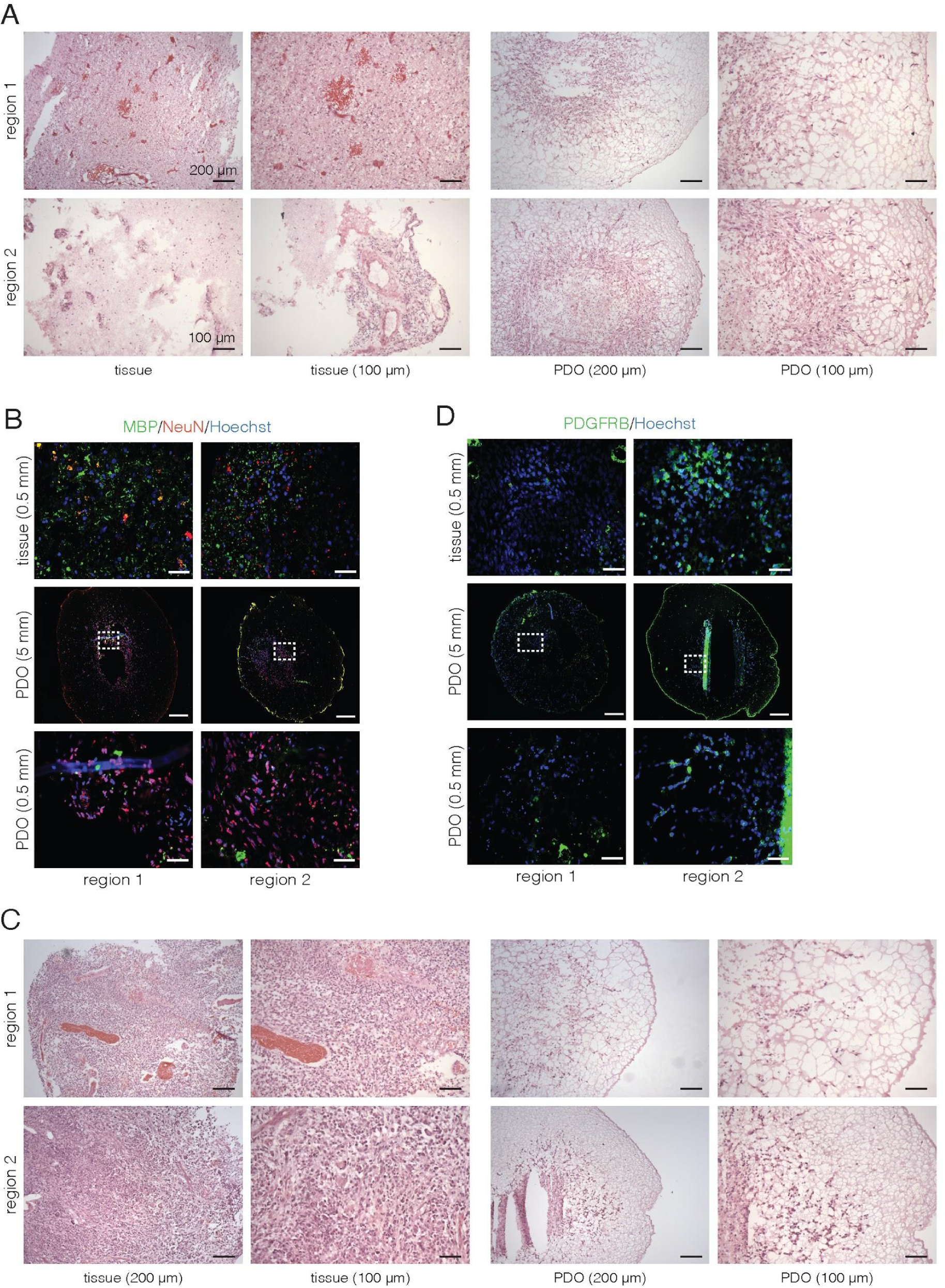
Histology and IF staining indicate higher proportions of malignant cells than scRNA-seq in select samples. **A** H&E staining of JK142 tissues and PDOs from both regions. The reg2 tissue sample appears to be largely composed of white matter infiltrated by a small population of neoplastic cells. Scale bar lengths indicated below the images. **B** IF images showing variable levels of MBP staining within and across JK142 tissue samples. White boxes on PDOs indicate the area shown in the zoomed-in images below. Scale bar lengths are indicated on the images’ left. **C** H&E staining of JK124 tissues and PDOs from both regions. The reg2 PDO appears largely composed of neoplastic cells rather than fibroblast-like cells as indicated by the scRNA-seq data. **D** IF images showing comparable levels of PDGFRB staining in reg1 and reg2 JK124 PDOs. White boxes on PDOs indicate the area shown in the zoomed-in images below.

**Supplementary Figure 6:**
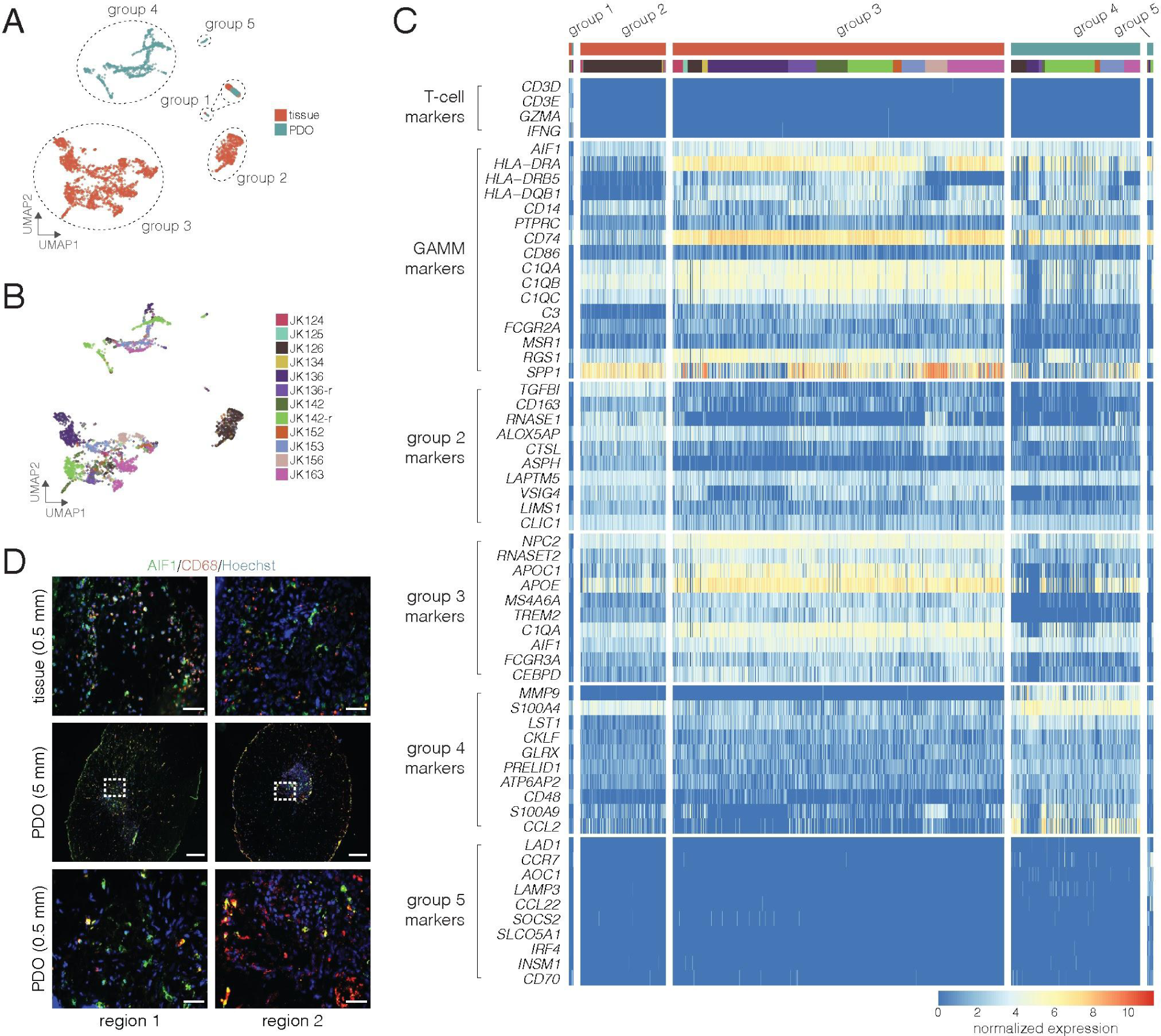
Macrophages found in PDOs are distinct from those found in matched tissue samples. **A-B** UMAP visualization of all tissue and PDO immune cells colored by sample type (A) and patient (B). Cell groups are circled and labeled in A. **C** Gene expression heatmap of select markers across immune cell groups. Both tissue and PDO cells are present in group 1 and express markers of cytotoxic T-cells. Cells in groups 2-4 variably express GAMM markers (Muller et al., 2017); the top 10 markers specific to each of these groups (relative to each other, see Methods) are also shown. **D** IF images showing the presence of GAMMs in tissues and PDOs (JK163). White boxes on PDOs indicate the area shown in the zoomed-in images below. Scale bar lengths are indicated on the images’ left.

**Supplementary Figure 7:**
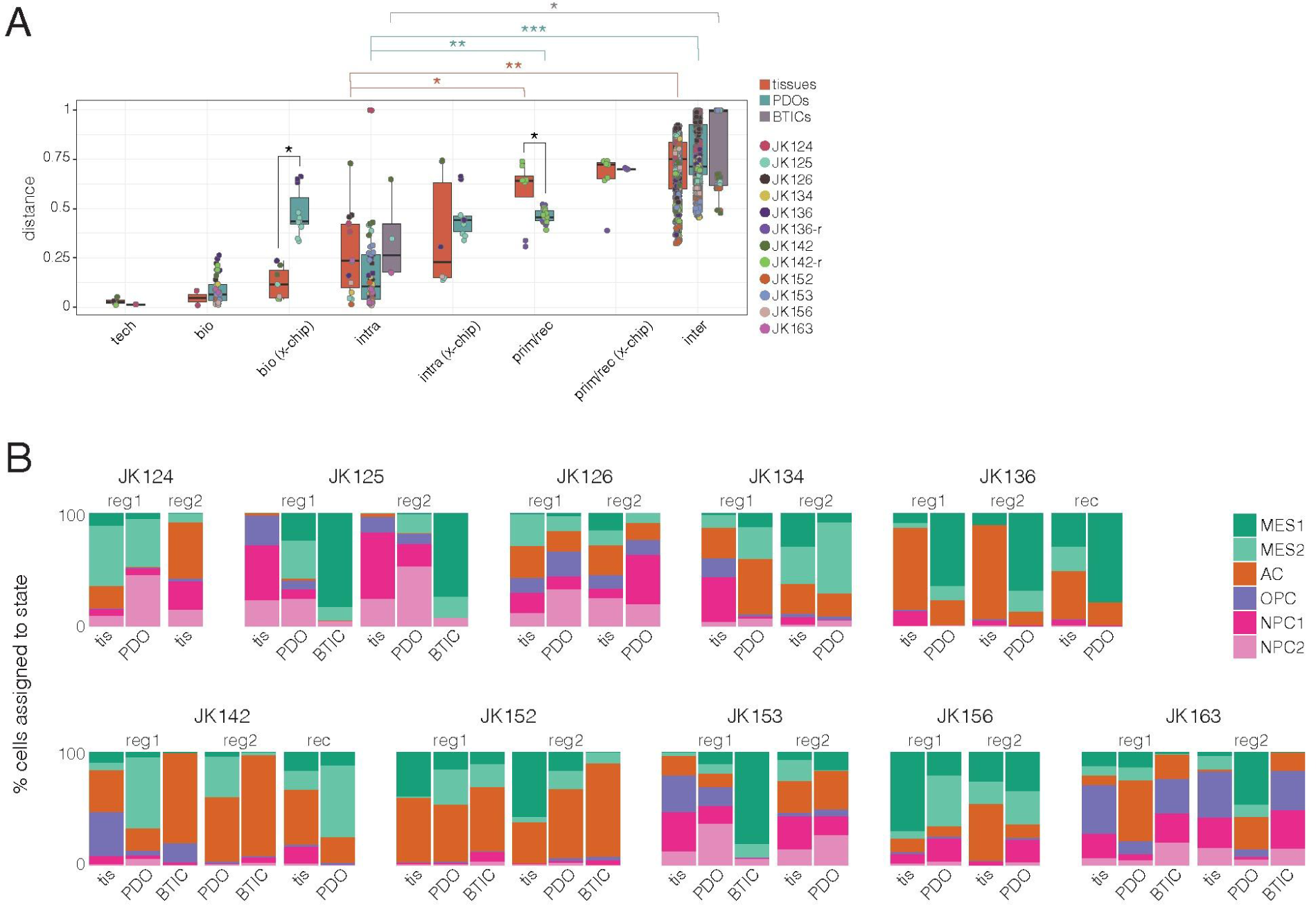
PDOs and BTICs variably retain characteristics of their parent tumors. **A** scUniFrac distances between different replicate types across tissue, PDO, and BTIC samples. Tech: same cell suspension run on different channels of the same 10X chip; bio: distinct tissue pieces or PDOs from the same tumor region; intra: samples from different regions of the same tumor; prim/rec: samples from matched primary and recurrent tumors; inter: samples from different patients; x-chip: indicated sample pair types processed in independent experiments on separate 10X chips. \**p* < 0.05, \*\**p* < 5×10^-9^, \*\*\**p* < 5×10^-16^ (Wilcoxon tests). **B** Proportions of cells assigned to indicated cell states. Cells were assigned to the state for which they scored highest (Methods).

## Supplementary Tables

**Supplementary Table 1: Clinical patient information and details of samples profiled by bWES, scWGS, and scRNA-seq.** Metrics summaries for scWGS and scRNA-seq datasets (output by the CellRanger pipeline) are also included.

**Supplementary Table 2: CNAs and SNVs/indels in GBM-associated genes across samples profiled by bWES.** Details of data shown in Figure 1D. CNAs were identified using TITAN, and SNVs/indels were identified using Strelka2 and Mutect2 (see Methods for details). Cellular prevalence reflects the estimated proportion of malignant cells carrying the event (*i.e.* a cellular prevalence <1 indicates a subclonal event).

**Supplementary Table 3:** Median exposure fractions of reference mutation signatures across samples profiled by bWES.

**Supplementary Table 4: Marker gene analysis results for tissue cell clusters.** Output of *overlapEpxrs* function from *scran* (see Methods for details). Top: the minimum rank of that gene across all pairwise comparisons. P.value: combined p-value across all pairwise comparisons, consolidated using Sime’s method. FDR: Benjamini-Hochberg-corrected p-values. Overlap proportions are defined as the probability that a randomly selected cell in the cluster of interest has a greater expression value than a randomly selected cell in the comparison cluster (indicated after “overlap”).

**Supplementary Table 5: Marker gene analysis results for PDO cell clusters.** Results as described for Supplementary Table 4.

**Supplementary Table 6: Marker gene analysis results for BTIC clusters.** Results as described for Supplementary Table 4.

**Supplementary Table 7: Marker gene analysis results for immune cell groups.** Results as described for Supplementary Table 4. The group 1 results are compared to every other group, while results for groups 2-5 were obtained from comparisons between these groups and excluding group 1.

**Supplementary Table 8: Genes over- or under-expressed in malignant PDO cells compared to malignant tissue cells.** Results are shown for each tumor/region pair, except JK124 reg2 and JK142 reg2, where only a single malignant PDO and tissue cell was identified, respectively.

**Supplementary Table 9: Genes over- or under-expressed in BTICs compared to malignant tissue cells.** Results are shown for each tumor/region pair where a BTIC line was profiled.

**Supplementary Table 10: Pathway enrichment analysis results for genes over- or under-expressed in malignant PDO cells or BTICs compared to malignant tissue cells.** The top 500 genes for every comparison (Supplementary Tables 8-9) were used as input for multi-sample enrichment analyses.

## Methods

### RESOURCE AVAILABILITY

#### Lead Contact and materials availability

Further information and requests for resources and reagents should be directed to and will be fulfilled by the Lead Contact, Marco Marra. There are restrictions to the availability of organoids due to the lack of an external centralized repository and our need to maintain the stock; however, organoids generated in this study will be made available upon reasonable request following approval by an internal review board and completion of a Materials Transfer Agreement.

#### Data and code availability

The datasets generated during this study will be made available upon publication. Code used to produce analyses and figures presented in this study can be found at https://github.com/vleblanc/GBM-PDO-paper.

### EXPERIMENTAL MODEL AND SUBJECT DETAILS

#### Human tissue samples

The use of tumor tissues and peripheral blood samples was coordinated through the Clark H. Smith Tumour and Related Tissue Bank. Tissues and blood samples were collected at the Foothills Medical Centre in Calgary, AB, Canada after informed patient consent under a protocol approved by the Health Research Ethics Board of Alberta Cancer Committee (HREBA.CC-16-0762). Additional work was done under HREBA.CC-16-0716 and a protocol approved by the UBC BC Cancer Research Ethics Board (H19-030103A005). All patient samples were de-identified prior to processing. Materials in this study were obtained from 10 patients (12 tumors) whose clinical information is summarized in Supplementary Table 1. The clinical molecular characterization included in Supplementary Table 1 was performed according to standard protocols in the Department of Neuropathology, Foothills Medical Centre, University of Calgary. Mutations were identified using an Agena (San Diego, CA, USA) MassARRAY custom molecular neuro panel. *MGMT* promoter methylation was assayed using methylation restriction enzyme digestion followed by real-time quantitative polymerase chain reaction (MRSE-qPCR), using the *OneStep* qMethyl^TM^-*Lite* kit from Zymo Research (Irvine, CA, USA) and custom *MGMT* promoter-specific primers/probes. A sample was considered to be methylated if the methylation score was greater than 9% (Dunn et al., 2009; Quillien et al., 2014).

### METHOD DETAILS

#### Collection and processing of patient GBM samples

All tissue was collected fresh at the time of surgical resection by the neurosurgeon (J.J.K.). Intra-operative frozen section was performed at the Department of Neuropathology (Foothills Medical Centre and University of Calgary) to confirm the diagnosis of GBM or to confirm the presence of active tumor in recurrent cases. Tissue for the study was then selected from the contrast-enhancing rim of the tumor by the neurosurgeon under direct visualization. Tissue from two distinct regions at opposite ends of each newly diagnosed tumor were obtained. For the two recurrent tumors, tissue was obtained from the contrast-enhancing area of the tumor. Tissue was carefully selected to maximize the amount of active tumor, and surrounding necrotic tissue or normal-appearing brain was removed directly by the surgeon. Tissue was then placed in sterile saline and taken immediately to the laboratory for processing.

In the laboratory, tissue was further subdivided into small pieces and processed for each component of the study. Tissue was processed immediately for organoid generation under sterile conditions in a laminar flow biosafety tissue culture hood (see below). Concurrent with organoid establishment, a number of tumor pieces were either flash-frozen in individual tubes by immersing in liquid nitrogen and then stored at -80°C or cryopreserved in 10% (v/v) DMSO by first incubating the samples in freezing solution at room temperature for 30 min and then transferring to -80°C in a CoolCell freezing container (Sigma-Aldrich, Oakville, ON, Canada). Additional tumor pieces were immediately fixed in 4% paraformaldehyde for histological and immunohistochemical studies. Tissue was fixed for approximately 1 hour at room temperature and subsequently cryoprotected by incubation in 30% sucrose (Sigma-Aldrich) at 4°C. Tumor pieces were then placed in a plastic mold and quickly frozen in OCT tissue freezing medium on dry ice. All frozen tissue was stored at -80°C in preparation for further processing.

#### Generation of patient-derived organoids and BTIC lines

To create Matrigel forms in which to grow the PDOs, we first placed strips of Parafilm (PM996, Fisher Scientific, Ottawa, ON, Canada) ∼1 in wide and 3-4 in long into a 15 cm petri dish (CA25383-103, VWR, Edmonton, AB, Canada) and created spherical depressions ∼2.5-3 mm in diameter to create a mold for the Matrigel. The forms were sterilized with 70% ethanol spray and stored at -20°C. To establish the PDOs, Matrigel HC (CABD354262, VWR) was thawed overnight at 4°C on ice and then, continuing to work on ice, 10 µL of thawed Matrigel was added to each Parafilm depression. Freshly resected tumor tissue was rinsed with sterile PBS and cut into small sections (∼1 mm^3^), then individual pieces were placed into the prepared Matrigel forms. The tumor-bearing Matrigel forms were allowed to solidify at room temperature for 10 min, after which a second drop (10 µL) of Matrigel was added to seal the tumor section within the Matrigel. The Matrigel forms were then placed in a 37°C incubator for 1 hour to promote Matrigel solidification, and then removed from the Parafilm and transferred to wells of a 24-well plate (one form per well) containing 1.5 mL of neural stem cell media (NeuroCult^TM^; 05700, STEMCELL Technologies, Vancouver, BC, Canada) supplemented with human EGF (20 ng/mL; AF-100-15, Peprotech, Rocky Hill, NJ, USA), FGF-2 (20 ng/mL; AF-100-18B, Peprotech), and heparin sulfate (2 µg/mL; 07980, STEMCELL Technologies). The plates were kept in a 37°C, 5% CO_2_, and 90% humidity sterile incubator and organoid growth was typically observed after 24-48 hours. Media was changed or added every 2-3 days until the cells originating from the tumor sections had grown out into the surrounding Matrigel (ranging from ∼20 days - 7 weeks), at which point the organoids were either (1) cryopreserved in 10% (v/v) DMSO by first incubating the samples in freezing solution at room temperature for 30 min and then transferring to -80°C in a CoolCell freezing container (Sigma-Aldrich), (2) fast-frozen in individual tubes by immersing in liquid nitrogen and then stored at -80°C, (3) embedded in an optimal cutting temperature (OCT) block and stored at -80°C, or (4) passaged by cutting the organoid into smaller pieces (typically 10) and re-implanting these into fresh Matrigel as described above.

In some cases, we observed cells escaping the Matrigel and creating neurospheres within the well. When this occurred, these cells were collected and propagated in 25 cm flasks under identical conditions as described above.

#### Immunostaining and imaging

Slides with cryosectioned tumor and organoid samples (7 μm) were dipped in acetone and dried overnight, followed by three 10 min PBS washes. Next, the slides were incubated in 300 µl of blocking solution (20% goat serum in PBT [10% normal goat serum/PBS with 0.3% Triton-X 100]) and their respective primary antibody for two hours at 37°C, followed by three 10 min PBS washes. The slides were then incubated with the appropriate fluorescently conjugated secondary antibody diluted in PBS (1/500) for one hour at 37°C, followed by three 10 min PBS washes. Hoechst nuclear stain was applied for one minute, followed by three 5 min PBS washes, and then slides were mounted. A Zeiss Axioplan 2 manual compound microscope with a Zeiss Axiocam HRc camera was used to capture the images. Adobe Photoshop was used to merge images acquired on different channels. Primary antibodies employed were as follows: Rabbit anti-Ki67 (Vector Laboratories; 1:300); Mouse anti-Nestin (STEMCELL Technologies; 1:100); Rabbit anti-GFAP (STEMCELL Technologies; 1:500); Mouse anti-Beta Tubulin (STEMCELL Technologies; 1:500); Rabbit anti-Olig2 (PhosphoSolutions; 1:300) Mouse anti-Sox2 (R&D Systems; 1:100), Goat anti-PDGFRa (R&D Systems; 1:100); Mouse anti-EGFR (Millipore; 1:100); Rabbit anti-Doublecortin (Abcam; 1:250); Mouse anti-Carbonic Anhydrase (CA9) (Abcam; 1:500), Rabbit anti-Myelin Basic Protein (BMP) (ThermoFisher; 1:300), Mouse anti-NeuN (Abcam; 1:500), Rabbit anti-CD45 (ThermoFisher; 1:200), Mouse anti-CD3 (ThermoFisher; 1:200), Rabbit anti-IBA1 (ThermoFisher; 1:200), Mouse anti-CD68 (ThermoFisher; 1:100), Rabbit anti-VIM (RayBiotech; 1:200), Mouse anti-Alpha-Smooth Muscle Actin (ACTA2) (ThermoFisher; 1:100), Rabbit anti-PDGFR beta (Abcam; 1:100) and Mouse anti-CD31 (ThermoFisher; 1:100). Secondary antibodies were Goat and Donkey anti-IgG and usually either Alexa Fluor-488, -555 or -568 conjugated (ThermoFisher Scientific; 1:500). All samples were counterstained with Hoechst nuclear stain (ThermoFisher Scientific; 1:2000).

#### Single-cell RNA-seq (scRNA-seq)

##### Dissociation into single-cell suspensions

Cryopreserved tissues or PDOs were rapidly thawed in a 37°C water bath. Tissues were washed once in NeuroCult NS-A Basal Medium (human; STEMCELL Technologies), and PDOs were washed in the same media at room temperature for 15 min with rocking. Samples were resuspended in 0.2 mg/mL collagenase (Sigma-Aldrich) plus 0.5 mg/mL DNaseI (Worthington Biochemical Corp, Lakewood, NJ, USA) and incubated at 37°C for 30 min with shaking every 10 min. Tissues were dissociated by tituration, incubated for another 10 min at 37°C, and filtered through a 40 μm mesh filter (BD Falcon, Bedford, MA, USA) with added NeuroCult^TM^ medium (STEMCELL Technologies) to remove larger pieces of debris and undissociated tissue. PDOs were dissociated by tituration and incubated for another 10 min at 37°C. Dissociated cells were then washed twice in Hank’s balanced salt solution (HBSS; Thermo Fisher Scientific, Waltham, MA, USA) and resuspended in phosphate-buffered saline (PBS; Thermo Fisher Scientific) with 0.04% (w/v) bovine serum albumin (BSA; Thermo Fisher Scientific). Additional HBSS washes were performed in samples with high debris and/or Matrigel content (up to four washes). Single-cell suspensions were filtered through 40 μm FlowMi tip filters (Bel-Art SP Scienceware, South Wayne, NJ, USA), and final cell concentrations were determined using the Countess automated cell counter (Thermo Fisher Scientific).

##### scRNA-seq library construction and sequencing

Single-cell 3’ RNA-seq libraries were generated using the Chromium Single Cell 3’ Library & Gel Bead Kit v2 (10X Genomics, Pleasanton, CA, USA) following the manufacturer’s protocol. Briefly, the provided targeted cell recovery table was used to determine the volume of cell suspension added to the Single Cell Master Mix prior to loading onto the Single Cell A chip (2,000 cells for tissue and PDO samples and 1,000 cells for BTIC lines). Following gel beads in emulsion (GEM) generation on the Chromium Controller, the GEM-RT reaction was performed, the GEMs were broken, and cDNA cleanup was performed using Dynabeads MyOne Silane beads. Samples that underwent evident clogs or wetting failures at the GEM generation step were not further processed. cDNA amplification was performed with 12 amplification cycles, and reaction cleanup was then performed using the SPRIselect reagent kit (Beckman Coulter, Indianapolis, IN, USA). Total cDNA quantification was performed using the Agilent High Sensitivity DNA Kit (Santa Clara, CA, USA). Enzymatic fragmentation, library construction, and additional SPRIselect clean-ups were then performed to generate the final indexed libraries. The number of sample index PCR cycles was determined based on the total cDNA yield and ranged from 10-14 cycles. Library quality was assessed using the Agilent High Sensitivity DNA kit and libraries were quantified using the Qubit dsDNA HS Assay kit (Thermo Fisher Scientific). Libraries were pooled (seven to eight libraries/pool) and sequenced on one lane of an Illumina HiSeq2500 flow cell (San Diego, CA, USA) using v3 or v4 chemistry and paired-end sequencing with single indexing following Illumina protocols and 10X sequencing parameters (26 bp read 1, 98 bp read 2, and 8 bp i7 index).

#### Single-cell whole-genome sequencing (scWGS)

##### Dissociation into single-nuclei suspensions

Nuclei isolation was performed following the Isolation of Nuclei for Single Cell DNA Sequencing Demonstrated Protocol (CG000167) from 10X Genomics. Briefly, a small piece (∼3-5 mm^3^) of flash-frozen tissue was cut from larger samples and immediately placed into a 1.5ml Eppendorf tube. Ice-cold lysis buffer (10mM Tris-HCl, 10mM NaCl, 3mM MgCl_2_, and 0.1% NP-40) was added and the tissue homogenized through an 18-gauge syringe followed by a 20-gauge syringe. The sample was briefly centrifuged (300 rcf) and the nuclei-containing supernatant was transferred to a fresh Eppendorf tube. The supernatant was further centrifuged (850 rcf) for 5 min to pellet the nuclei, and the nuclei were washed twice and then resuspended in ice-cold Nuclei Wash & Resuspension buffer (0.04% [w/v] BSA in PBS). Final nuclei concentrations were determined by DAPI staining and counting using the Countess automated cell counter or through manual hemocytometer counts. Cryopreserved organoids were directly placed into 1.5ml Eppendorf tubes and lysed as per tissue sections (4-5 organoids combined).

##### scWGS library construction and sequencing

Single-cell DNA libraries were generated using the Chromium Single Cell DNA Reagent Kit (10X Genomics) following the manufacturer’s protocol. Briefly, targeted cell recovery (250 cells) was used to determine the volume of cell suspension added to the Single Cell Bead Mix prior to loading onto the Chromium Chip C alongside the CB polymer. The generated cell beads were then shaken on a thermomixer overnight. The encapsulated cells were then lysed and the genomic DNA (gDNA) was denatured. The reaction mix, cell beads (containing the denatured DNA), and gel beads were loaded onto the Chromium D Chip to generate GEMs on the Chromium Controller. Following GEM generation, an isothermal incubation was performed, the GEMs were broken, and DNA cleanup was performed using Dynabeads MyOne Silane beads followed by a SPRIselect reaction cleanup. Total DNA quantification was performed using the Agilent High Sensitivity DNA Kit. Library construction and additional SPRIselect clean-ups were then performed to generate the final indexed libraries, using 12 sample index PCR cycles. Library quality was assessed using the Agilent High Sensitivity DNA kit and libraries were quantified using the Qubit dsDNA HS Assay kit. All libraries were pooled together and initially sequenced on four lanes of an Illumina HiSeqX flow cell using v2.5 chemistry and paired-end sequencing with single indexing following Illumina protocols and 10X sequencing parameters (100 bp read 1, 100 bp read 2, and 8 bp i7 index). Following preliminary analysis of recovered cells per sample, libraries were pooled and further sequenced to reach a median depth of ∼750,000 reads/cell.

##### Whole-exome sequencing (bWES)

Isolated nuclei (as described in *Single-cell whole-genome sequencing*), flash frozen tissue sections (cut into small pieces), cryopreserved organoids, or peripheral blood from matched patients were used as sample sources for genomic DNA, which was isolated using the Qiagen DNEasy Blood and Tissue Kit (Toronto, ON, Canada) or the Qiagen QIAamp DNA Mini Kit following the manufacturer’s instructions. Total DNA was quantified using the Qubit dsDNA HS Assay kit.

Genomic DNA libraries were constructed according to Canada’s Michael Smith Genome Sciences Centre plate-based and paired-end library protocols on a Microlab NIMBUS liquid handling robot (Hamilton, USA). Briefly, 1-100 ng of total nucleic acid was sonicated (Covaris LE220) in 62 µL volume to 250-350 bp. Sonicated DNA was purified with PCRClean DX magnetic beads (Aline Biosciences). The DNA fragments were end-repaired, phosphorylated and bead-purified in preparation for A-tailing using a custom NEB Paired-End Sample Prep Premix Kit (New England Biolabs). Illumina sequencing adapters were ligated for 15 minutes at 20°C and adapter-ligated products were then bead-purified and enriched with 10 cycles of PCR using primers containing a hexamer index that enables library pooling.

Five or six different libraries (total of >500 ng) were pooled prior to whole-exome capture using the xGen® Exome Research Panel v1.0 (Integrated DNA Technologies). The pooled libraries were hybridized to the capture probes at 65°C for a minimum of four hours. Following hybridization, streptavidin-coated magnetic beads (Dynabeads M-270) were used for exome capture, and then post-capture enrichment was performed using six PCR cycles and primers that maintain the library-specific indices. The pooled libraries were sequenced with paired-end 125 base reads in two lanes of an Illumina HiSeq2500 flow cell using v4 chemistry.

### QUANTIFICATION AND STATISTICAL ANALYSIS

#### Preprocessing of 10X scWGS data

##### Count matrix generation

The CellRanger-dna software (v1.1) was used to create fastq files (cellranger-dna mkfastq with default parameters) and to perform alignment, filtering, barcode counting, coverage analysis, and CNV calling (cellranger-dna cnv). The CellRanger-dna GRCh38-1.0.0 reference was used for alignment.

##### Data preprocessing

Data preprocessing was performed in R using the imputed CNV calls output by CellRanger-dna. Cells with at least 50 effective reads per Mb were retained for further analysis. To minimize the effect of noise, we calculated bin sizes that would have on average 200 reads/bin across samples (as recommended by 10X) for each patient and took the mean copy number across these coarse-grained bins.

##### Analysis of scWGS data

Clones were identified as described in (Laks et al., 2019). Coarse-grained copy number profiles were first reduced using UMAP (McInnes et al., 2018) with the *umap* package in R (v0.2.2.0; manhattan distance and num_neighbors = 15) and then clusters were identified using HDBSCAN (*dbscan* R package, v1.1-5) (Hahsler et al., 2019). Cells were over-clustered (*minPts = 10*) and then clusters were merged if their median copy number profiles overlapped by at least 90% (JK142 and JK153) or 80% (JK136; copy number resolution was lower in this sample due to increased ploidy in a substantial fraction of cells). Merged clusters were designated as non-malignant if the cluster was composed of diploid cells lacking large-scale copy number variants, otherwise as malignant clones. Unassigned cells and clusters composed of visibly noisy profiles were removed.

##### Preprocessing of bWES data

Following the removal of reads that failed Chastity filtering (Kircher et al., 2011), raw reads were aligned to the human reference genome GRCh38/hg38 with alternate contigs removed (no alt; https://www.bcgsc.ca/downloads/genomes/9606/hg38_no_alt/bwa_0.7.6a_ind/genome/) using the Burrows-Wheeler Aligner (BWA-MEM, v0.76a) (Li, 2013). BAM files were sorted and reads were merged and marked for duplicates using sambamba (v0.5.5).

#### Analysis of bWES data

##### SNV/indel calling

To identify somatic mutations in tissue and organoid samples, SNVs and indels were called using both Strelka2 and Mutect2 and the intersection of variants identified using these two approaches were considered as true positives (Cai et al., 2016). For the Strelka2 approach, Manta (v1.6.0) was first run to identify candidate small indels, which were then passed to Strelka2 (v2.9.10) (Kim et al., 2018). In both cases, the matched blood sample was used as a normal comparator, the *--exome* flag was used, and the target regions were provided. SnpSift (from SnpEff v4.1) (Cingolani et al., 2012) was used to retain variants that passed the empirical variant scoring (EVS) filter. The union of variants identified in all samples from a single patient (tissues and PDOs) was then obtained, using RTG Tools (v3.10.1) to identify common variants.

Mutations were then also called using Mutect2 (Benjamin et al., 2019) from GATK (v4.1.4.1). The multisample mode was used to call SNVs/indels from all samples from each patient (tissues and PDOs), with the matched blood sample again used as a normal comparator. Exome target regions were provided, and the gnomAD exome r2.2.1 (GRCh38 liftover) variants (Karczewski et al., 2020) were used as a germline resource, as recommended by the GATK best practices. Variants that passed the *filterMutectCalls* (run with default parameters) filter were retained for downstream analyses.

The intersect of variants identified by each of these methods (obtained using RTG Tools) was considered for downstream analyses. Variants were annotated for their effect using SnpEff (GRCh38.79) and for SNPs (dbSNP v149) using SnpSift.

##### Copy number calling

Copy number profiles were obtained using Titan CNA (Ha et al., 2014). The snakemake workflow (https://github.com/gavinha/TitanCNA/tree/master/scripts/snakemake) was used, with R3.6 and setting *genomeBuild* to hg38. Default configuration settings were otherwise used as suggested by the package authors.

##### Variant clustering and subclone identification

In order to identify groups of variants with similar characteristics across patient-specific samples, we used PyClone to cluster variants identified from all samples for each patient, and obtain the cellular prevalence of these clusters within individual samples (Roth et al., 2014). To create the PyClone input tables, we used the final variants identified for each patient group as discussed above (using the allele read depths output by Mutect2 output) and the Titan CNA output. Variants were filtered to have a total read depth >= 20 and an alternate read depth >= 10. Due to the large number of variants identified in the JK163 reg2 PDO sample, an alternate read depth filter of 50 was used for this sample. Variants in regions with subclonal copy number alterations (*i.e.* with cellular prevalence > 0.25 or < 0.75, as smaller or larger subclones are less likely to impact variant alternate read depths) were also removed.

We then used CITUP (Malikic et al., 2015) to identify clonal populations and their phylogenetic relationships, as described by McPherson et al. (2018). Specifically, for each patient the *cluster.tsv* file file output by PyClone was reshaped into a table of mean cellular prevalence values, with clusters as rows and samples as columns, and then the iterative version of CITUP was run on this table. If the number of clusters output by PyClone was larger than eight, the *--max-nodes* argument was set to this number in order to allow for the optimal solution to contain one node per cluster; otherwise, default parameters were used. The optimal tree solution was obtained from the hdf5 results file (/results/optimal), and its clonal phylogeny and clone frequencies obtained from the /trees/{tree_solution}/adjacency_list entry and the /trees/{tree_solution}/clone_freq entry, respectively. Plotting of the optimal tree solution was inspired by the *mapscape* R package; the *igraph* R package (v1.2.5) was used to plot the tree, with the edge lengths modified to be scaled to the number of variants gained in each clone.

##### Mutation signature analysis

To try to understand the mechanisms underlying mutational profiles in different samples, we performed mutation signature analysis using the SignIT R package (Zhao et al., 2017) and the Catalog of Somatic Mutations in Cancer (COSMIC) single base substitution (SBS) signatures (v3) (Alexandrov et al., 2020) as reference.

#### Combined analysis of scWGS and bWES data

To investigate the relationships between mutation-based and copy number-based clonal populations, we analyzed the scWGS and bWES data together. For each copy number clone, we first created new BAM files containing only reads from cells assigned to that clone: to do this, we first used the *cellranger-dna bamslice* function to create clone-specific BAM files containing only reads from cells assigned to that clone within each sample (*e.g.* “clone a” cells in reg1 tissue, clone a cells in reg1 PDO, etc.), and then merged these using *sambamba* (v0.6.1) across all samples from a single patient (*e.g.* “clone a” files from each sample belonging to a single patient, etc.) Each of these patient- and clone-specific BAM files were then queried for variants identified in the patient of interest (see the *SNV/indel calling* section) using the *mpileup* (with the *--ignore-RG* flag) and *call* (with the *-m* and *-Oz* flags) functions from *bcftools* (v1.9).

#### Preprocessing of 10X scRNA-seq data

##### Count matrix generation

The CellRanger software from 10X Genomics was used to create fastq files (cellranger mkfastq with default parameters; v2.0.0 for all samples except those from JK163 and the “batch” samples, for which v2.1.1 was used) and to perform alignment, filtering, barcode counting, and UMI counting (cellranger count with default parameters except --expect-cells set to 1000 for all samples; v2.1.1). The CellRanger GRCh38-1.2.0 reference was used for alignment.

##### Data preprocessing and normalization

Data preprocessing was performed in R, based on the filtered count matrices output by CellRanger. For each sample, outlier cells were identified as those with > 3 median absolute deviations (MADs) below the median total UMI counts or total number of genes detected, or with > 3 MADs above the median percent of counts from mitochondrial genes (Lun et al., 2016b), and were discarded (5,046/86,532 cells, 5.8%). Counts for remaining cells from all samples (81,486 cells) were combined into a single count matrix, and genes detected (UMI > 0) in at least 10 cells and with at least 20 UMIs across all cells were retained. In subsequent analyses, any time the matrix was subset these settings were applied to identify retained genes unless otherwise specified. Normalization was then applied to all cells using the *scran* package in R (Lun et al., 2016a). The *quickCluster* function was used to cluster cells for normalization (with min.mean = 0.1 as suggested for UMI data), and the resulting clusters were used as input to the *computeSumFactors* function (with min.mean = 0.1). These factors were then used in the *normalize* function of the *scater* R package.

To identify putative doublets, we applied the *doubletCells* function from the *scran* package to each sample, with k = 30. Cells with outlier doublet scores (> 3 MADs above the sample median) were discarded (6,411/81,486 cells, 7.9%), and normalization was performed again on the resulting dataset. The final matrix thus consisted of 75,075 cells and 23,076 genes that were used in downstream analyses. Unless otherwise specified, normalized expression values were used for all analyses.

#### Analysis of scRNA-seq data

##### Cell clustering

Cell clustering was performed largely as described in Shekhar *et al*. (2016). Highly variable genes (HVGs) were first identified using the *makeTechTrend* function of the *scran* R package, assuming Poisson-distributed technical noise. This was done individually for each tumor, and the variance decompositions were combined using the *combineVar* from *scran*, using the weighted “z” method. The *decomposeVar* function was then used to decompose gene-specific variances into biological and technical components, and genes with a biological component > 0.01 and a Benjamini-Hochberg-corrected *p*-value < 0.05 were considered HVGs. Principal component analysis (PCA) was performed using the *multiBatchPCA* function from *scran*, considering only HVGs and setting patient ID as batch. The Horn’s parallel analysis approach implemented in the *parallelPCA* function from *scran* was used to identify significant principal components (PCs), with the exception that the permuted iterations (100) were also run using *multiBatchPCA*. To perform Louvain-Jaccard clustering (Levine et al., 2015; Shekhar et al., 2016), we first created a *k*-nearest neighbor graph using the *nn2* function of the *RANN* R package (*k* = 30 for most analyses, *k* = 15 for immune cell clustering analyses). Edges in the graph were then weighed by the Jaccard overlap index of the neighborhood of corresponding nodes. Finally, the weighted graph was used as input for Louvain clustering (Blondel et al., 2008), performed using the *igraph* R package.

To explore relationships between the clusters identified, we used a method analogous to the one used in the *groupDistance* function of the *Seurat* R package. Specifically, mean PC values were first calculated across significant PCs for each cluster. A distance matrix was built using Euclidean distance weighted by the relative contribution of each PC’s eigenvalue and used as input for hierarchical clustering using the complete linkage method (using the *hclust* function in R). This tree was also used to calculate generalized UniFrac distances using code from the *scUnifrac* package (v0.9.6) (Liu et al., 2018) modified to use a previously calculated tree.

##### Visualization

UMAP was performed using the *umap* R package, based on PCA space using only significant PCs as described above. The number of neighbors was set to the same value of *k* used for clustering.

##### Comparisons to reference cell types

Cells were scored according to their similarity to reference cell types using the *SingleR* R package (v0.2.0) (Aran et al., 2019) with the default references. The HPCA (Human Primary Cell Atlas) main reference was used in the figures shown in Supplementary Figure 3.

##### Differential gene expression analyses

DE analyses were performed using the *overlapExprs* function from *scran*, with default arguments except *direction = “up”* to only identify over-expressed genes in each comparison. Genes labeled as feature controls were not included. In the cluster-specific DE analyses, tumor identity was used as a blocking factor. For the DE analysis performed between immune cell groups, the combined tumor identity and sample type (tissue or PDO) was used as a blocking factor. For the DE analysis performed between sample types (*e.g.* tissue vs PDO cells), individual analyses were performed for each tumor/region pair.

##### Copy number inference

Copy number was inferred from scRNA-seq data using the infercnv tool (v1.1.1) from the Trinity CTAT project (https://github.com/broadinstitute/inferCNV). For the tissue cells, bulk RNA-seq data from normal brain samples was first used as a reference as previously described (Patel et al., 2014). Data were obtained from the Genotype-Tissue Expression (GTEx) project (https://gtexportal.org/home/datasets): “Brain - Cerebellum”, “Brain - Caudate (basal ganglia)”, “Brain - Cortex”, “Brain - Nucleus accumbens (basal ganglia)”, “Brain - Cerebellar Hemisphere”, “Brain - Frontal Cortex (BA9)”, and “Brain - Hippocampus” samples were used (gene TPM files downloaded on June 22, 2019). Cells that were consistently identified as non-malignant based on expression clustering results and on clustering of inferred copy number profiles were then used as a reference to re-calculate copy number profiles in malignant cells. Immune cells were excluded from this new reference due to their unique expression profile in chromosome 6 (Supplementary Figure 4). The non-malignant tissue cells were also used as references for calculating inferred copy number in PDO cells and BTICs. For all analyses, the “denoise” argument was set to TRUE and a minimum average expression cutoff of 0.1 was used, and was calculated exclusively in the single-cell data (*i.e.* bulk GTEx samples were excluded).

##### Enrichment analyses

Enrichment analyses were performed on the top 500 over-expressed and the top 500 under-expressed genes identified in each regional pair or primary/recurrent pair using Metascape (Zhou et al., 2019) based on the following gene sets: GO Biological Processes, Reactome Gene Sets, Hallmark Gene Sets, and Oncogenic Signatures. Results were extracted from the “_FINAL_GO.csv” file, and only the top term for each of the top 10 groups are shown in Figure 4.

##### Cell state signatures

Cell state scores and two-dimensional coordinates were calculated as described in (Neftel et al., 2019). Cells were assigned to the state or meta-state (quadrant) for which they scored highest. Differential enrichment within individual quadrants between tissue and PDO cells or tissue cells and BTICs was calculated using Fisher’s exact test on the proportion of cells from each source/region that were present in the quadrant of interest. Shannon’s index for each sample was calculated based on the number of malignant cells assigned to each cell state using the *entropy* function from the *entropy* R package (v1.2.1; with *method = “ML”*).

